# mTORC1 Directs TNF-Induced Life-or-Death Decisions via Complex I Destabilization

**DOI:** 10.64898/2026.06.28.735146

**Authors:** Zhao Deng, Jingyi Wang, Buhao Deng, Kai Yu, Yan Liu, Jinglin Xu, Haiwen Lin, Jiafan Yuan, Tingyun Yang, Haotian Wang, Bo Yan, Youwei Ai

## Abstract

The transition from tumor necrosis factor α (TNF)-induced plasma membrane-bound complex I to cytosolic death-inducing complex II switches cells from survival to death. However, the precise regulation of this fatal decision is incompletely understood. Here, we show that mTORC1 promotes this transition by destabilizing later-stage complex I without affecting its initial assembly. Inhibition of mTORC1 unleashes ATG9A and FIP200 activity, thereby promoting the accumulation of CHUK (IKKα) in complex I. CHUK scaffolds the kinase-active IKKβ to stabilize complex I and prevent complex II formation. Activation of this ATG9A/FIP200–CHUK/IKKβ axis protects against TNF-induced fulminant hepatitis while compromises antibacterial defense against *Staphylococcus aureus*. This mTORC1-governed life-or-death transition provides therapeutic insight into TNF-related pathologies—including cancer, metabolic tissue injury, and microbial infections—where mTORC1 activity is frequently suppressed.

## INTRODUCTION

Tumor necrosis factor α (TNF) is a pleiotropic cytokine that orchestrates diverse cell fate decisions through its receptor TNFR1. Under basal conditions, TNF predominantly activates pro-survival signaling pathways, including NF-κB and MAPKs. However, when one or more intracellular cell death checkpoints are disrupted, TNF stimulation can shift toward the induction of cell death.^1,2^ Disruption of these checkpoints—through agents such as interferon γ (IFNγ),^3–5^ Smac mimetics (Smac),^6^ transcriptional inhibitors, or translational inhibitor cycloheximide (CHX) ^7^—triggers cell death in various cell types in the presence of TNF. Upon stimulation, TNF engages TNFR1 to initiate the assembly of a membrane-bound early complex I within minutes, composed of death domain (DD)-containing adaptors RIPK1 and TRADD. This complex I recruits E3 ligases (TRAF2, cIAP1/2, and the LUBAC complex: HOIL1, HOIP, SHARPIN) and kinases (TAK1, TBK1, and CHUK(IKKα)/IKKβ), which induce site-specific ubiquitination and phosphorylation of multiple complex I components.^1,8–10^ TNF-induced cell death is initiated only after RIPK1 and/or TRADD dissociate from membrane-bound complex I to assemble a cytosolic complex II by recruiting FADD and Caspase-8, leading to Caspase-8 activation.^7,11^

The transition from membrane-bound complex I—following a period of signal propagation—to cytosolic complex II represents a central step in TNF-induced cell death.^11^ However, the mechanisms driving this transition remain incompletely understood. In this study, we uncover a multilayered regulatory mechanism governed by mTORC1 that unexpectedly destabilizes later-stage complex I, thereby enabling the formation of complex II and promoting cell death. Given that mTORC1 activity is frequently altered in tumors, metabolic tissue injury, and infections,^12–16^ our findings uncover a new axis for therapeutic intervention in TNF-related pathologies.

## RESULTS

### The Enzymatic Activity of mTORC1 Promotes TNF-Induced Apoptosis

Proteins that mediate the transition from complex I to complex II are thought to be essential for TNF-induced cell death, and their inhibition is expected to prevent cell death. To identify factors involved in the transition, we performed a genome-wide CRISPR-Cas9 screen in HeLa cells, selecting for gene knockouts that conferred resistance to TNF combined with IFNγ, Smac, or CHX treatment (Figure 1A). In parallel, a targeted small-molecule screen of U.S. FDA-approved compounds was conducted to identify inhibitors of TNF/IFNγ-induced apoptosis (Figure S1A). The CRISPR screen enriched gRNAs targeting known apoptotic regulators—Caspase-8, FADD, and Bid—as well as unexpected components of the mTORC1 complex, including mTOR, the scaffold Raptor, and Rag-Ragulator subunits LAMTOR1 and LAMTOR4 (Figures 1B and 1C). Supporting these genetic findings, the compound screen identified 12 structurally distinct mTOR enzymatic inhibitors that suppressed TNF-induced cell death (Figures S1B and S1C). Together, these results suggest that mTORC1 enzymatic activity is required for TNF-induced apoptosis.

**Figure 1.**
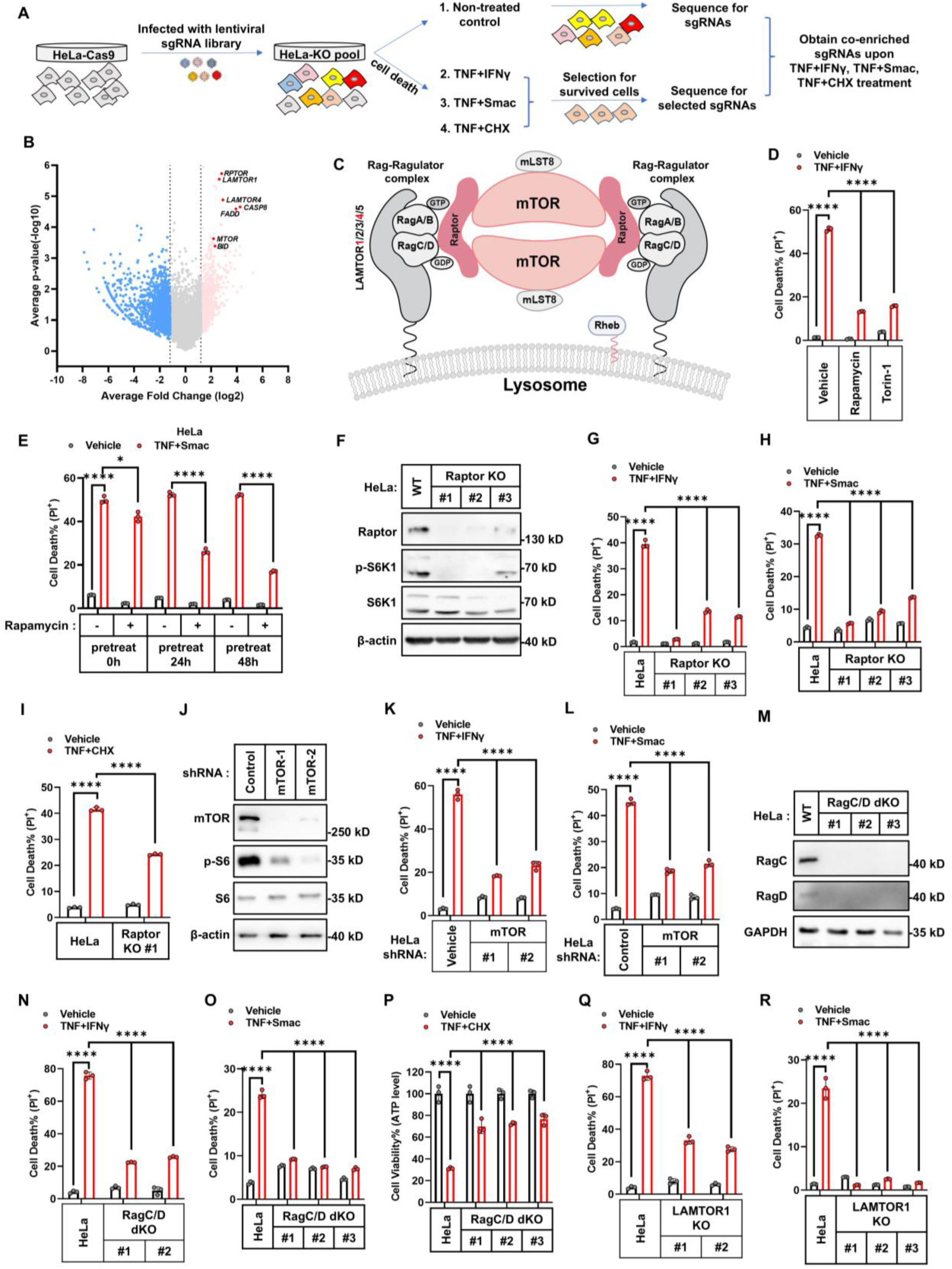
mTORC1 promotes TNF-induced cell death. (**A**) Schematic of the CRISPR-Cas9 screening strategy used to identify shared regulators involved in TNF+IFNγ-, TNF+Smac-, and TNF+CHX-induced cell death. (**B**) Representative volcano plot showing Log_2_-transformed average fold change of sgRNA abundance in HeLa-Cas9 cells treated with TNF combinations versus control. Genes significantly enriched under all conditions are labeled. (**C**) Illustration of the mTORC1 complex localized at the lysosomal surface via the Rag-Ragulator complex. (**D**) HeLa cells were treated with TNF+IFNγ for 48 h in the presence or absence of mTOR kinase inhibitors rapamycin (100 nM) or Torin1 (100 nM), and cell death was measured. **(E)** HeLa cells were pretreated with rapamycin for 0 h, 24 h, or 48 h, followed by TNF+Smac treatment and cell death was assessed. **(F)** Raptor KO in HeLa cells (clones #1, #2, and #3 generated using three different gRNAs) was confirmed by immunoblotting. (**G-I**) HeLa cells (parental, Raptor KO clones #1, #2, #3) were treated with TNF+IFNγ (G), TNF+Smac (H) for 48h, or TNF+CHX for 8h (I). Cell death was measured by PI staining. **(J)** HeLa cells stably expressing control or mTOR shRNA (#1 or #2) were analyzed by immunoblotting for mTOR expression. (**K, L**) HeLa cells stably expressing control shRNA or mTOR shRNA (#1 or #2) were treated with TNF+IFNγ (K) or TNF+Smac (L), and cell death was measured. **(M)** RagC/D dKO in HeLa cells (clones#1, #2 and #3) was confirmed by immunoblotting. (**N-P**) HeLa cells (parental, RagC/D dKO clones#1, #2 and #3) were treated with TNF+IFNγ (N), TNF+Smac (O) or TNF+CHX (P), and cell death was measured. (**Q, R**) HeLa cells (parental, LAMTOR1 KO clones #1 and #2) were treated with TNF+IFNγ (Q), or TNF+Smac (R), and cell death was measured. Vehicle was used as control for specific stimuli. Cell death in D, E, G, H, I, K, L, N, O, Q, R was measured by PI staining followed by flow cytometry (n=3). Cell death in P was assessed using ATP-based luminescence assay. The data are shown as mean ± SD, with n = 3 biologically independent samples. Statistical significance in all panels was evaluated using two-way ANOVA: ****P < 0.0001; *P=0.03 (E); ns, not significant.

To validate this, we treated various human and murine cancer cell lines, a normal human hepatocyte line (HL-7702), and primary mouse bone marrow–derived macrophages (BMDMs) with two mechanistically distinct mTORC1 inhibitors—rapamycin and Torin-1.^17^ Both inhibitors significantly reduced TNF/IFNγ- and TNF/Smac-induced cell death, as measured by propidium iodide (PI) staining and ATP levels (see below) (Figures 1D, 1E, S1D–S1M), but had no effect on TRAIL-induced apoptosis, even after 48-hour of rapamycin pretreatment (Figures S1N and S1O). Raptor knockout in HeLa, HCT116, and MCF7 cells (Figures 1F-1I, S2A–S2F), or in primary mouse hepatocytes (Figures S2G and S2H), abolished TNF/IFNγ-, TNF/Smac-, and TNF/CHX-induced apoptosis. Reconstitution with wild-type Raptor restored mTORC1 signaling (indicated by S6K1 or S6 phosphorylation ^18^) and cell death sensitivity, whereas a Rag-binding–deficient Raptor mutant failed to do so (Figures S2I-S2L). Similarly, knockdown or knockout of mTORC1 components—mTOR, RagC and RagD, or LAMTOR1—suppressed TNF-induced apoptosis (Figures 1J-1R, S2M–S2O), while Rictor knockout (mTORC2-specific) had no effect (Figures S2P–S2W).

The Rag-Ragulator complex—but not mTORC1’s enzymatic activity—has been reported to mediate Caspase-8–dependent GSDMD cleavage and pyroptosis during *Yersinia* infection. ^19^ To determine whether it is the catalytic activity of mTORC1, rather than the scaffolding function of the Rag-Ragulator complex, that drives TNF-induced apoptosis, we designed a system in which mTORC1 activity is preserved despite disruption of the Rag-Ragulator complex. This was achieved by leveraging the mechanism whereby the lysosome-localized Rag-Ragulator complex activates mTORC1 through recruitment of Raptor to the lysosomal membrane ^20^ (Figure 2A). By fusing Raptor to a lysosomal localization signal, we restored mTORC1 activity in RagC/D double knockout or LAMTOR1 knockout cells (Figures 2B, 2C, 2D, 2G and 2I). Restoration of mTORC1 activity in these Rag-Ragulator–deficient cells reinstated TNF-induced cell death (Figures 2E, 2F, 2H, 2J and 2K). Together, these findings demonstrate that the enzymatic activity of mTORC1, rather than Rag-Ragulator–mediated scaffolding, is essential for TNF-induced apoptosis—a mechanism distinct from that observed during *Yersinia*-induced pyroptosis.

**Figure 2.**
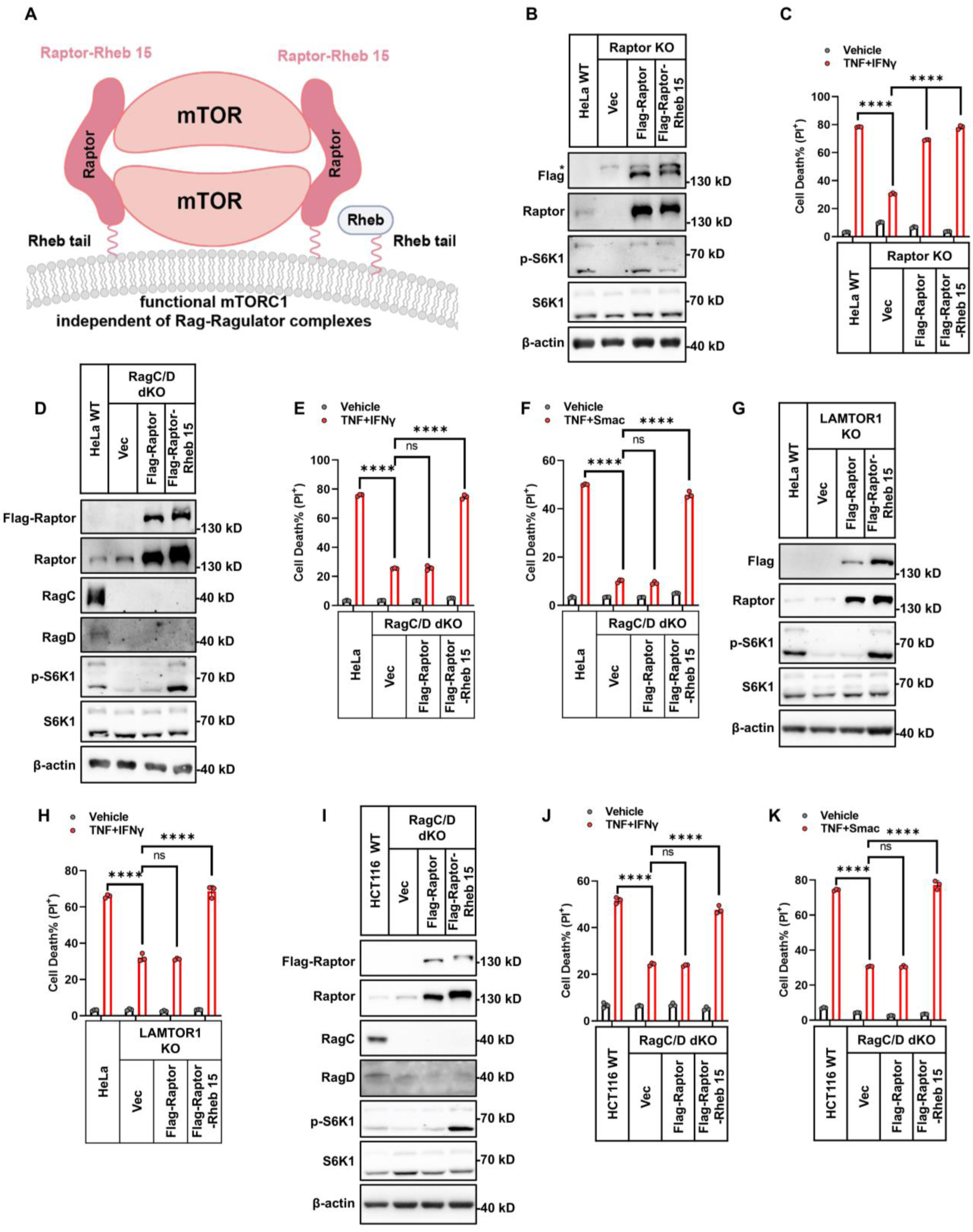
mTORC1 Kinase Activity Drives TNF-Induced Cell Death. (**A**) Schematic of the functional mTOR-Raptor-Rheb15 complex in Rag-Ragulator-deficient background. **(B)** Immunoblot analysis of Raptor and p-S6K1 level in HeLa cells (parental, Raptor KO, and Raptor KO cells stably expressing Flag-Raptor or Flag-Raptor-Rheb15). Asterisk indicates non-specific bands. **(C)** The indicated HeLa cells were treated with TNF+IFNγ for 48 h. Cell death was measured by PI staining followed by flow cytometry (n = 3). **(D)** Immunoblot analysis of Raptor, RagC, RagD, and p-S6K1 in HeLa cells (parental, RagC/D dKO, and RagC/D dKO cells reconstituted with Flag-Raptor or Flag-Raptor-Rheb15). (**E**, **F**) HeLa cells (parental, RagC/D dKO, and RagC/D dKO stably expressing Flag-Raptor or Flag-Raptor-Rheb15) were treated with TNF+IFNγ (E) or TNF+Smac (F), and cell death was measured. **(G)** Immunoblot analysis of Raptor and p-S6K1 in HeLa cells (parental, LAMTOR1 KO, and LAMTOR1 KO cells reconstituted with Flag-Raptor or Flag-Raptor-Rheb15). (**H**) HeLa cells (parental, LAMTOR1 KO, and LAMTOR1 KO reconstituted with Flag-Raptor or Flag-Raptor-Rheb15) were treated with TNF+IFNγ, and cell death was measured. **(I)** Immunoblot analysis of Raptor, RagC, RagD, and p-S6K1 in HCT116 cells (parental, RagC/D dKO, and RagC/D dKO cells reconstituted with Flag-Raptor or Flag-Raptor-Rheb15). **(J, K)** The indicated HCT116 cells were treated with TNF+IFNγ (J) or TNF+Smac (K) for 48 h. Cell death was measured by PI staining followed by flow cytometry (n = 3). Vehicle was used as control for specific stimuli. The data are shown as mean ± SD, with n = 3 biologically independent samples. Statistical significance in all panels was evaluated using two-way ANOVA: ****P < 0.0001; ns, not significant.

### mTORC1 Drives Complex I to Complex II Transition via Destabilization of Later-Stage Complex I

Since mTORC1 inhibition blocks TNF-induced apoptosis, we investigated which step—complex I disassembly, complex II formation, or Caspase-8 activation—is disrupted upon mTORC1 suppression. We first assessed Caspase-8 activation. Both pharmacological and genetic inhibition of mTORC1 reduced the levels of cleaved Caspase-8, Caspase-3, and PARP (Figures 3A, 3B, S3A–S3F). This reduction was not due to enhanced lysosomal degradation (Figures S3G and S3H) or extracellular secretion of Caspase-8 (Figure S3I).

**Figure 3.**
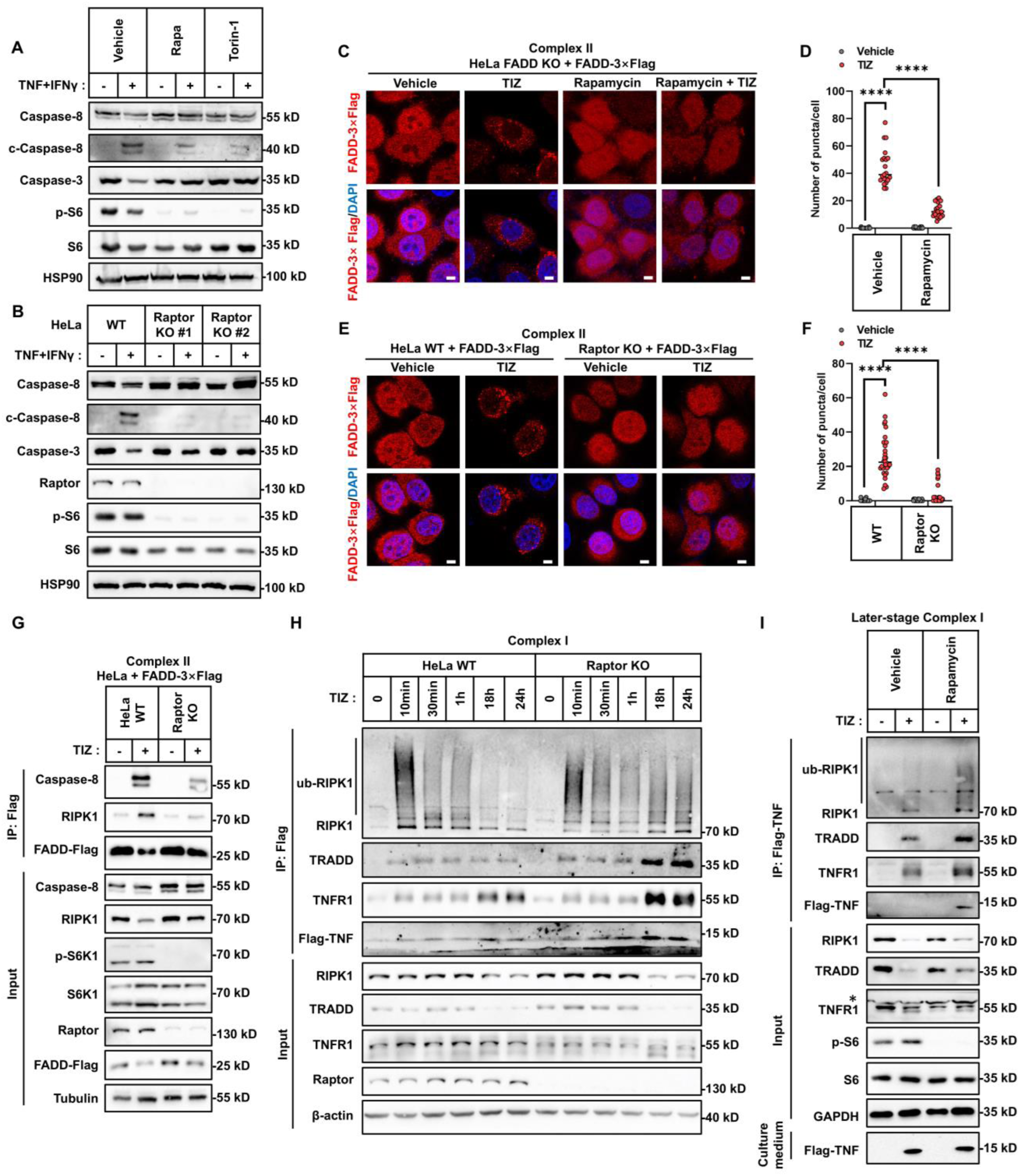
mTORC1 facilitates the transition of complex I to complex II in TNF-induced cell death. (**A**) HeLa cells were treated with TNF+IFNγ for 36 h in the presence or absence of rapamycin or Torin-1. The expression of cleaved and full-length Caspase-8 and caspase-3 was analyzed by immunoblotting. (**B**) HeLa cells (parental and Raptor KO clones #1 and #2) were treated with TNF+IFNγ for 36 h. The expression of cleaved and full-length Caspase-8 and Caspase-3 was analyzed by immunoblotting. (**C**, **D**) Representative confocal images (C) of FADD KO HeLa cells stably expressing FADD-3×Flag treated with TNF+IFNγ+z-VAD (TIZ) for 24 h, with or without rapamycin. Cells were co-stained with anti-Flag antibody and DAPI. Cytosolic complex II was visualized as FADD-Flag puncta. Quantification of puncta per cell is shown in (D). Scale bar: 5 μm, n=21 cells. (**E**, **F**) Representative confocal images (E) of parental or Raptor KO HeLa cells stably expressing FADD-3×Flag treated with TIZ for 24 h. Cells were co-stained with anti-Flag antibody and DAPI. Cytosolic complex II was visualized as FADD-Flag puncta. Quantification is shown in (F). Scale bar: 5 μm, n=21 cells. (**G**) HeLa cells (WT or Raptor KO) stably expressing FADD-3×Flag were treated with TIZ for 24 h. Cytosolic complex II was immunoprecipitated using anti-Flag antibody, and co-immunoprecipitated proteins were detected by immunoblotting. (**H**) HeLa cells (WT or Raptor KO) were treated with Flag-TNF+IFNγ+z-VAD (TIZ) for the indicated times. Membrane-bound complex I was immunoprecipitated using anti-Flag antibody and analyzed by immunoblotting. (**I**) HeLa cells were treated with Flag-TNF+IFNγ+z-VAD (TIZ) for 24 h in the presence or absence of rapamycin (100 nM). Complex I was analyzed by immunoblotting. The data are shown as mean ± SD, with n = 3 biologically independent samples. C and E show representative immunofluorescence images from two independent experiments. All immunoblots are representative of three independent experiments. Statistical significance in panels D and F was evaluated using the Kruskal–Wallis test: ****P < 0.0001; ns, not significant.

We next examined whether complex II formation is impaired under mTORC1 inhibition. Using microscopy to visualize C-terminally tagged FADD (3×Flag or GFP), we found that mTORC1 inhibition reduced the formation of cytosolic FADD or FADD–RIPK1 puncta in multiple cell types following TNF stimulation (Figures 3C–3F, S3J-S3W). This was further confirmed by co-immunoprecipitation of 3×Flag-tagged FADD, which showed reduced association with RIPK1 and Caspase-8 in both human and mouse cells (Figures 3G, S3X-S3A1). Notably, the pan-Caspase inhibitor z-VAD-FMK (z-VAD) was included to prevent rapid disassembly of complex II and enhance detection ^7,21^ (Figures S3M and S3Z).

Reducing complex I formation or enhancing NF-κB activity can potentially suppress complex II assembly and subsequent cell death. ^11,22^ However, inhibition of mTORC1 does not affect early complex I formation or the activation of NF-κB and MAPK pathways (Figures S3B1-S3D1). Since complex I assembles within minutes after TNF stimulation, while cell death typically occurs much later (TNF/IFNγ and TNF/Smac require ∼18 and ∼12 hours, respectively; TNF/CHX requires several hours), we assessed the stability of later-stage complex I at the onset of cell death or when cell death became evident.

Despite comparable early complex I formation (Figure 3H), mTORC1 inhibition markedly stabilized later-stage TNF–TNFR1–RIPK1–TRADD-containing complex I in both human and mouse cells (Figures 3H, 3I, S3E1-S3K1). z-VAD was used to inhibit cell death and preserve protein recovery for immunoprecipitation; it did not, on its own, affect complex I stability (Figure S3F1). Stabilization of RIPK1 and/or TRADD within complex I impedes the formation of cytosolic complex II and thus suppresses cell death. This is further supported by our observation that expression of a ubiquitination-deficient RIPK1 K377R ^23,24^ mutant in Raptor/RIPK1 double knockout cells, which destabilizes RIPK1 K377R within complex I, led to more pronounced cell death compared to cells expressing wild-type RIPK1 (Figures S3L1 and S3M1). Thus, mTORC1 facilitates the transition from complex I to complex II by destabilizing later-stage complex I, thereby enabling TNF-induced cell death.

### mTORC1 Destabilizes Later-Stage Complex I via ATG9A and FIP200

We hypothesized that mTORC1 inhibition activates proteins it normally suppresses, which in turn stabilize complex I and prevent cell death. If so, knocking out these proteins should restore cell death despite mTORC1 inhibition. To identify such factors, we conducted a CRISPR-Cas9 screen in Raptor knockout cells under TNF/IFNγ stimulation and searched for depleted gRNAs indicative of reactivated cell death (Figure S4A). This screen revealed significant loss of gRNAs targeting two autophagy proteins, ATG9A and FIP200 (Figure 4A), suggesting their knockout restores sensitivity to cell death in mTORC1-inhibited cells.

**Figure 4.**
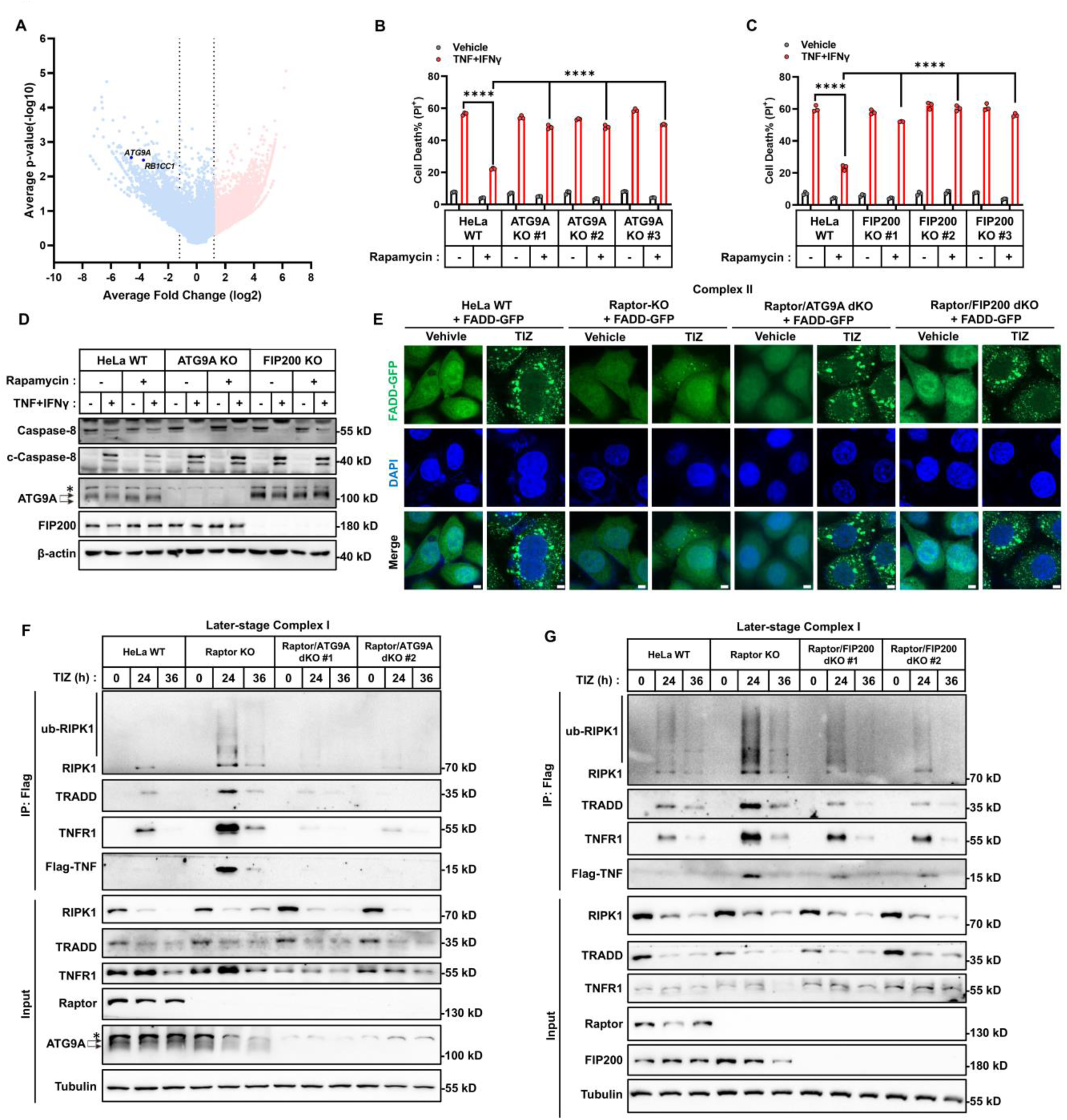
Stabilization of later-stage complex I requires ATG9A and FIP200. (**A**) HeLa Raptor KO-Cas9 cells were transduced with a genome-wide CRISPR-Cas9 gRNA library and treated with TNF+IFNγ for 36 h, followed by DNA isolation and deep sequencing. A volcano plot shows average fold change of sgRNA abundance in TNF+IFNγ-treated versus control cells. ATG9A and RB1CC1 (FIP200) are labeled. (**B**) HeLa cells (parental and ATG9A KO clones #1, #2, #3) were treated with TNF+IFNγ in the presence or absence of rapamycin (100 nM) for 48 h. Cell death was measured by PI staining followed by flow cytometry (n = 3). (**C**) HeLa cells (parental and FIP200 KO clones #1, #2, #3) were treated as in (B), and cell death was measured (n = 3). (**D**) HeLa cells (parental, ATG9A KO, and FIP200 KO) were treated with TNF+IFN γ in the present or absent of rapamycin (100 nM) for 36 h. Cleaved and full-length Caspase-8 levels were analyzed by immunoblotting. Asterisk indicates non-specific band. (**E**) Representative confocal images of HeLa cells (parental, Raptor KO, Raptor/ATG9A dKO, and Raptor/FIP200 dKO), stably expressing FADD-GFP, treated with TIZ for 24 h. Cytosolic complex II formation was visualized as GFP-positive puncta. Nuclei were counterstained with DAPI. Scale bar, 5 μm. (**F**) HeLa cells (parental, Raptor KO, and Raptor/ATG9A dKO clones #1 and #2) were treated with Flag-TNF+IFNγ+z-VAD (TIZ) for the indicated times. Membrane-bound complex I was immunoprecipitated using anti-Flag antibody, and representative proteins were analyzed by immunoblotting. Asterisk indicates non-specific bands. (**G**) HeLa cells (parental, Raptor KO, and Raptor/FIP200 dKO clones #1 and #2) were treated as in (F), and membrane-bound complex I was analyzed. The data are shown as mean ± SD, with n = 3 biologically independent samples. Panel E shows representative immunofluorescence images from three independent experiments. All immunoblots are representative of two to three independent experiments. Statistical significance in all panels was evaluated using two-way ANOVA: ****P < 0.0001; ns, not significant.

Indeed, deletion of ATG9A or FIP200 increased cell death in mTORC1-deficient HeLa, HCT116, and MCF7 cells to levels comparable with mTORC1-proficient wild-type cells (Figures 4B, 4C, S4B-S4P), indicating that the protective effect of mTORC1 inhibition requires ATG9A/FIP200. Consistently, mTORC1 inhibition blocked complex II formation, Caspase-8 activation, and stabilized membrane-bound later-stage complex I in an ATG9A- and FIP200-dependent manner (Figures 4D–4G, S4Q-S4X, and later in Figures S7Q, S7R).

Our findings recall a recent study identifying a TAX1BP1–FIP200–ATG13–ATG9A-mediated noncanonical autophagy pathway that promotes lysosomal degradation of complex II and cleaved Caspase-8, thereby protecting cells from TNF-induced cell death.^25^ In that work, knockout of TAX1BP1, FIP200, ATG13, or ATG9A rendered cells sensitive to otherwise nonlethal doses of TNF.^25^ In contrast, our model differs in several key aspects: (i) In the prior study, ATG9A/FIP200 suppressed cell death under nonlethal TNF stimulation in the presence of active mTORC1, whereas in our system—under lethal TNF triggers—ATG9A/FIP200 inhibit cell death only when mTORC1 is inactive; (ii) ATG9A and FIP200 stabilize later-stage complex I rather than mediate complex II degradation; (iii) cleaved Caspase-8 is not degraded via the lysosome upon mTORC1 inhibition (Figures S3G and S3H).

Furthermore, knockout of TAX1BP1 or ATG13 did not phenocopy ATG9A or FIP200 loss in our system (Figures S4Y-S4C1), indicating that cell protection upon mTORC1 inhibition is not due to TAX1BP1–FIP200–ATG13–ATG9A-mediated complex II degradation. Notably, in mouse EMT6 cells, ATG9A knockout modestly but significantly sensitized cells to TNF/IFNγ- and TNF/Smac-induced death even when mTORC1 was active (Figures S4D1-S4F1, S4J1), whereas this effect was marginal in three human cell lines (Figures 4B, 4C, S4D, S4M, S4N, S4G1, S4I1). This suggests that noncanonical autophagy ^25^ and complex I stabilization may co-occur in a cell-type-specific manner, further supporting that these are distinct pathways. Importantly, ATG9A remains essential for the protective effect of mTORC1 inhibition in EMT6 cells (Figures S4E1, S4F1). In summary, mTORC1 inhibition–dependent stabilization of membrane-bound later-stage complex I requires ATG9A and FIP200, and the resulting blockade of cell death is not due to TAX1BP1–FIP200–ATG13–ATG9A-mediated complex II degradation.

### ATG9A/FIP200 promote CHUK/IKKβ accumulation to stabilize later-stage complex I upon mTORC1 inhibition

Since neither ATG9A nor FIP200 was detected in complex I by mass spectrometry or immunoprecipitation (see below), we hypothesized that downstream proteins regulated by ATG9A/FIP200 are translocated to complex I to mediate its stabilization during mTORC1 inhibition. To identify such candidates, we performed LC-MS/MS on complex I–associated proteins and cross-referenced those upregulated upon mTORC1 inhibition with the dropout hits (Figures 4A and S5A). CHUK (IKKα) and IKKβ emerged as top candidates (Figure S5A).

Knockout of CHUK or IKKβ, or pharmacological inhibition of both using TPCA-1,^26^ restored TNF-induced cell death in mTORC1-inhibited human and mouse cells (Figures 5A–5D, S5B-S5Q), indicating their non-redundant roles in mediating mTORC1-dependent protection. Although both are known NF-κB/MAPK activators, CHUK KO only modestly impaired these pathways, whereas IKKβ KO or TPCA-1 treatment caused stronger inhibition (Figures S6A-S6D), suggesting that CHUK/IKKβ regulate cell death in an NF-κB–independent manner when mTORC1 is inhibited.

**Figure 5.**
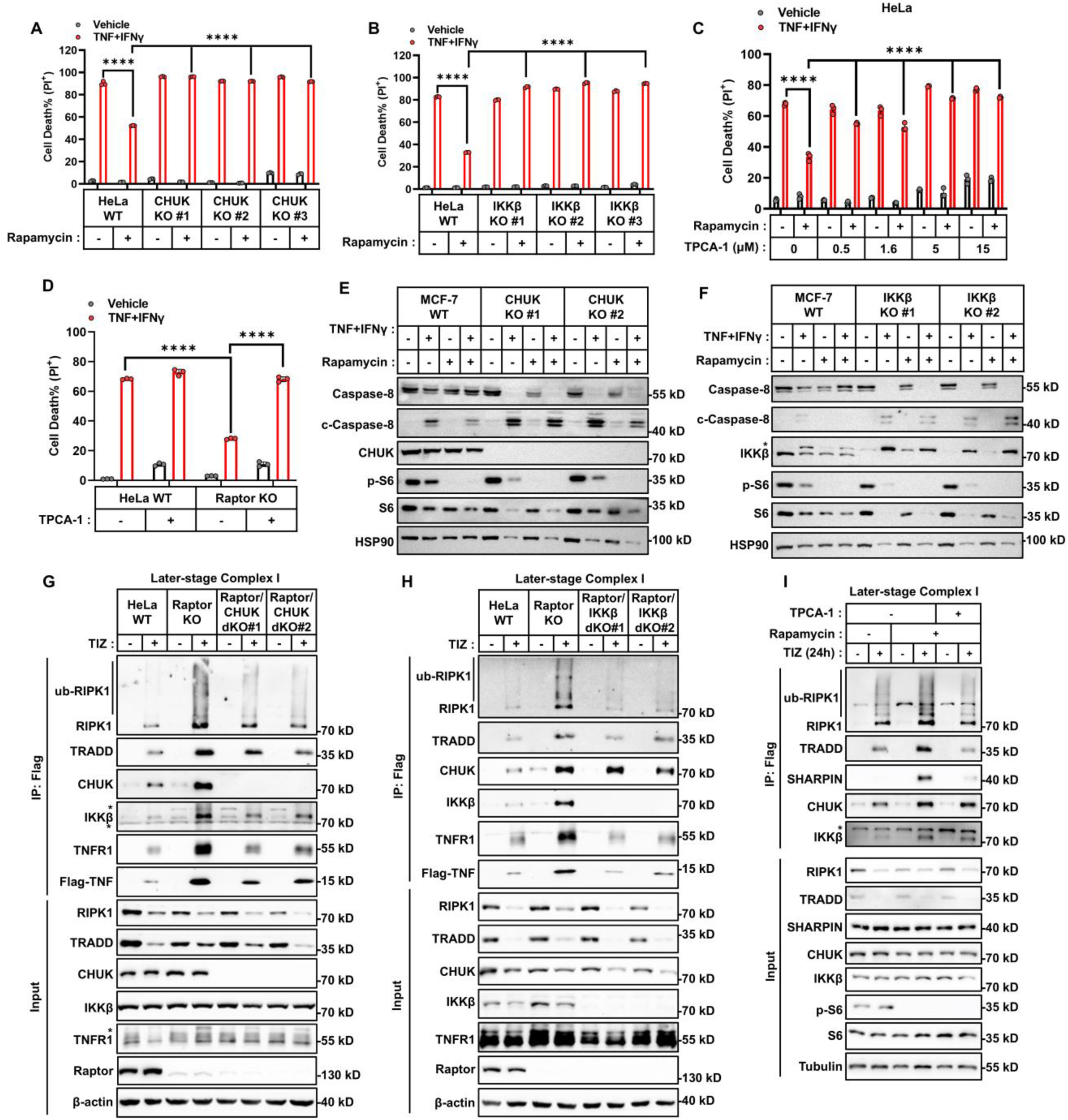
CHUK/IKKβ stabilizes later-stage complex I to restrict Caspase-8 activation. **(A**, **B**) HeLa cells (parental and CHUK (A) or IKKβ (B) KO clones #1, #2, #3) were treated with TNF+IFNγ in the presence or absence of rapamycin for 48 h. Cell death was measured by PI staining followed by flow cytometry (n=3). **(C)** HeLa cells were treated with TNF+IFNγ in the presence or absence of rapamycin with different concentrations of TPCA-1 for 48 h. Cell death was measured by PI staining followed by flow cytometry (n=3). **(D)** HeLa cells (parental, Raptor KO) were treated with TNF+IFNγ in the presence or absence of TPCA-1 (0.5μM) for 48 h. Cell death was measured (n = 3). **(E)** Immunoblot analysis of full-length and cleaved Caspase-8 in MCF-7 WT and CHUK KO cells treated with TNF+IFNγ with or without rapamycin (100 nM) for 36 h. **(F)** Immunoblot analysis of Caspase-8 and cleaved Caspase-8 in MCF-7 WT and IKKβ KO cells treated with TNF+IFNγ with or without rapamycin (100 nM) for 36 h. (**G**) HeLa cells (parental, Raptor KO, Raptor/CHUK dKO clones #1, 2) were treated with Flag-TNF+IFNγ+z-VAD (TIZ) for 24 h. Membrane-bound complex I was immunoprecipitated using anti-Flag antibody, and representative proteins were analyzed by immunoblotting. (**H**) HeLa cells (parental, Raptor KO, Raptor/IKKβ dKO clones #1, 2) were treated with TIZ for 24 h. Membrane-bound complex I was analyzed. (**I**) HeLa cells were treated with TIZ in the presence or absence of TPCA-1 or rapamycin for 24 h, and membrane-bound complex I was determined. The data are showed as mean ± SD, with n = 3 biologically independent samples. All representative immunoblots are from two to three independent experiments. Asterisk indicates non-specific bands. Statistical significance in all panels was evaluated using two-way ANOVA: ****P < 0.0001; ns, not significant.

As expected, mTORC1 inhibition enhanced the association of CHUK and IKKβ with complex I (Figures 5G and 5H). Disruption of CHUK or IKKβ, or treatment with TPCA-1, destabilized later-stage complex I and increased Caspase-8 activation (Figures 5E-5I, S5R-S5T), indicating that this association is required for the stabilization of complex I. Paradoxically, TPCA-1 impaired complex I stability without reducing associated CHUK or IKKβ levels (Figure 5I), suggesting that the accumulation of CHUK and IKKβ is a functional upstream event rather than a consequence of complex I stabilization.

To clarify the specific roles of CHUK and IKKβ in complex I stabilization, we examined their recruitment dynamics. CHUK knockout abolished mTORC1 inhibition–mediated association of IKKβ with later-stage complex I (Figures 5G and S5S). In contrast, IKKβ knockout had no effect on CHUK levels (Figures 5H and S5T), suggesting that CHUK is recruited first and subsequently facilitates the recruitment of IKKβ.

This points to a scaffolding role for CHUK, rather than a kinase-dependent function. Supporting this, re-expression of kinase-dead CHUK mutants in Raptor/CHUK double KO cells restored IKKβ recruitment, stabilized complex I, and suppressed cell death similarly to wild-type CHUK (Figures 6A–6C). In contrast, kinase-dead IKKβ failed to block cell death in mTORC1-inhibited cells (Figures 6D, 6E, S6E-S6G). Thus, CHUK and IKKβ function sequentially—CHUK as a scaffold and IKKβ as the essential kinase—rather than redundantly.

**Figure 6.**
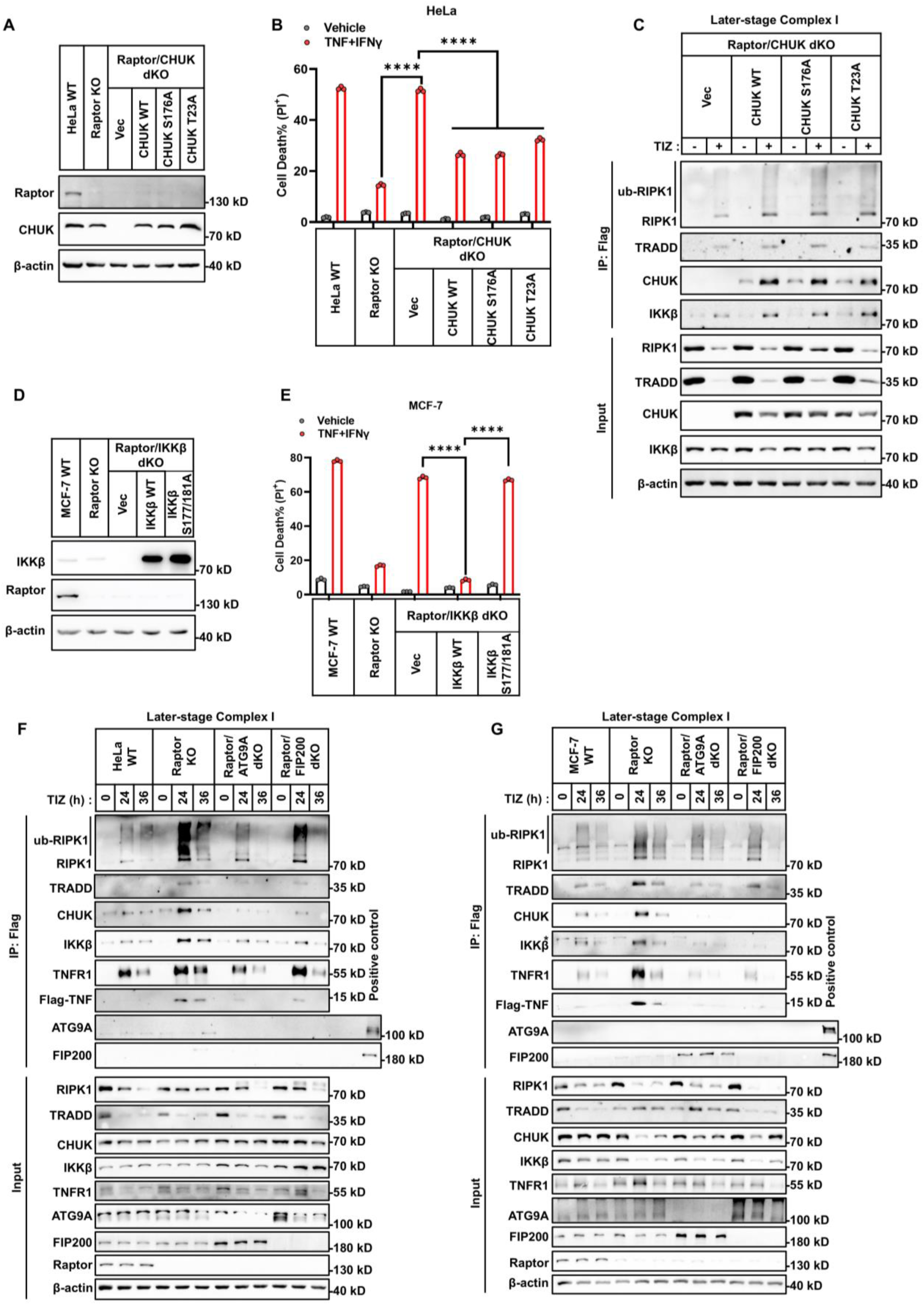
CHUK scaffolds kinase-active IKKβ to stabilize later-stage complex I. **(A)** Immunoblot analysis of Raptor and CHUK expression in HeLa cells (parental, Raptor KO, and Raptor/CHUK dKO). The Raptor/CHUK dKO cells were stably reconstituted with either an empty vector, WT CHUK, or kinase-deficient CHUK mutants (S176A or T23A). **(B)** The indicated HeLa cells were treated with TNF+IFNγ for 48 h. Cell death was assessed by PI staining and flow cytometry (n = 3). **(C)** HeLa Raptor/CHUK dKO cells reconstituted with vector, CHUK WT, S176A, or T23A mutants were treated with Flag-TNF+IFNγ+z-VAD (TIZ) for 24 h in the presence or absence of rapamycin. Membrane-bound complex I was isolated via Flag immunoprecipitation and representative proteins were analyzed by immunoblotting. **(D)** Immunoblot analysis of IKKβ expression in MCF-7 cells. The Raptor/IKKβ dKO cells were stably reconstituted with either an empty vector, WT IKKβ, or kinase-deficient IKKβ mutants (S1177A and S181A). **(E)** The indicated MCF-7 cells were treated with TNF+IFNγ for 48 h. Cell death was assessed by PI staining and flow cytometry (n = 3). **(F, G)** HeLa (F) or MCF-7 (G) cells (parental, Raptor KO, Raptor/ATG9A dKO, Raptor/FIP200 dKO) were treated with TIZ for 24 h or 36 h, and complex I was analyzed. The data are showed as mean ± SD, with n = 3 biologically independent samples. All representative immunoblots are from two to three independent experiments. Asterisk indicates non-specific bands. Statistical significance in all panels was evaluated using two-way ANOVA: ****P < 0.0001; ns, not significant.

Given that ATG9A and FIP200 are required for complex I stabilization, we tested whether they regulate CHUK/IKKβ accumulation. Although neither ATG9A nor FIP200 is incorporated into complex I, their knockout in Raptor KO cells abolished CHUK and IKKβ enrichment on later-stage complex I (Figures 6F and 6G). In summary, mTORC1 inhibition triggers an ATG9A/FIP200-dependent enrichment of CHUK to complex I, which in turn recruits IKKβ. This sequential assembly stabilizes complex I via IKKβ’s kinase activity, ultimately preventing TNF-induced cell death.

### mTORC1 inhibition activates the ATG9A/FIP200–CHUK/IKKβ axis to protect against TNF-induced hepatitis but impairs antibacterial immunity

To assess whether mTORC1 inhibition protects against TNF-mediated inflammatory tissue damage via stabilization of complex I by the ATG9A/FIP200–CHUK/IKKβ axis, we used a low-dose LPS/D-Galactosamine (LPS/D-Gal)–induced hepatitis model. D-Gal is metabolized specifically in the liver and inhibits hepatocyte transcription, rendering cells sensitive to TNF-induced apoptosis.^27–30^

LPS/D-Gal treatment induced Caspase-8 and Caspase-3 cleavage, resulting in cell death (TUNEL^+^), liver injury, and lethality in both male and female mice (Figures S7A, 7A-7F). Double knockout of Caspase-8 and RIPK3—but not RIPK3 alone—prevented lethality, confirming TNF-induced apoptosis as the main driver (Figures S7E). As a control, Concanavalin A (ConA)–induced autoimmune hepatitis, which depends on T cell- and macrophage-mediated injury,^31,32^ did not induce Caspase-8 cleavage, and Caspase-8 knockout failed to rescue survival (Figures S7B-S7D, S7F).

As expected, hepatocyte-specific Raptor deletion (Raptor^f/f^ × Alb-Cre) protected against LPS/D-Gal- but not ConA-induced liver damage and death, indicating that mTORC1 inhibition blocks TNF/Caspase-8–dependent injury (Figures S7A-S7D). In contrast, hepatocyte-specific ATG9A/Raptor double knockout restored sensitivity to LPS/D-Gal (Figures 7A–7F). Similarly, treatment with the CHUK/IKKβ kinase inhibitor TPCA-1 abolished protection conferred by Raptor deletion (Figure S7G).

**Figure 7.**
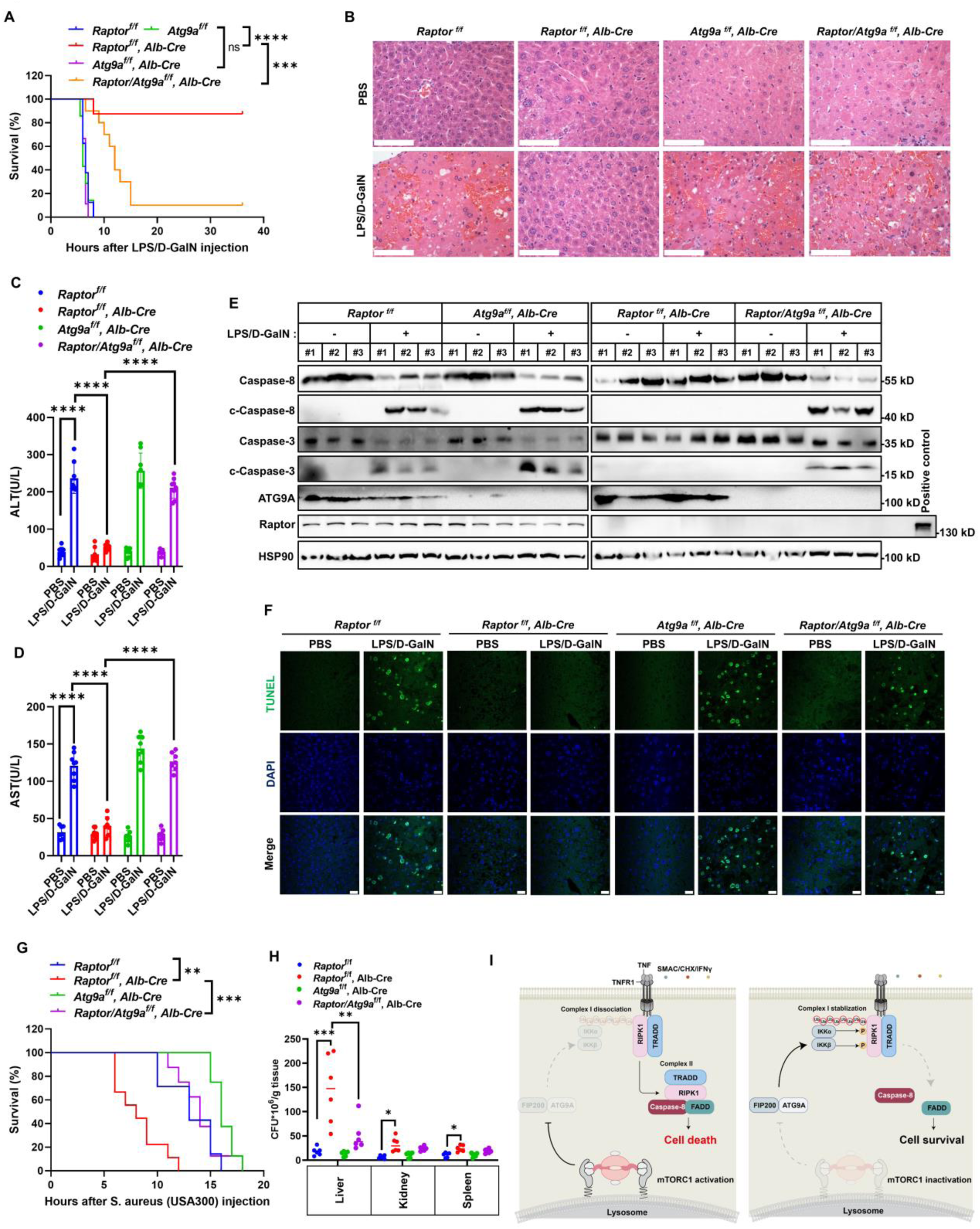
The ATG9A/FIP200-CHUK/IKKβ axis protects against TNF-induced hepatitis while impairs antibacterial immunity. (**A**) Cumulative survival curves of male C57BL/6J mice (8 weeks old) with indicated genotypes (*Raptor^fl/fl^, Atg9a^fl/fl^, Raptor^fl/fl^ Alb-Cre, Atg9a^fl/fl^ Alb-Cre, and Raptor/Atg9a^fl/fl^ Alb-Cre*) intraperitoneal injected with LPS (10 µg/kg) plus D-GalN (1 g/kg) (n = 10). (**B**) Representative hematoxylin and eosin (H&E) staining of liver sections from mice of the indicated genotypes 4 h after LPS/D-GalN challenge. Scale bar: 50 μm. (**C**, **D**) Serum levels of ALT (C) and AST (D) were measured 4 h after LPS/D-GalN injection (n = 6). (**E**) Immunoblot analysis of cleaved Caspase-8 and cleaved Caspase-3 in liver lysates from mice of the indicated genotypes (n = 3). (**F**) Representative images of TUNEL staining showing apoptotic cell death in liver sections 4 h post LPS/D-GalN challenge (n = 6). Scale bar: 20 μm. (**G**) Cumulative survival of male C57BL/6J mice with indicated genotypes (*Raptor^fl/fl^, Raptor^fl/fl^ Alb-Cre, Atg9a^fl/fl^ Alb-Cre,* and *Raptor/Atg9a^fl/fl^ Alb-Cre*) following intravenous injection of *S. aureus* USA300 (4×10^8^ CFU) (n = 8-10). (**H**) Bacterial burden (CFU/g tissue) in the liver, kidney, and spleen of mice 24 h after intravenous infection with *S. aureus* USA300 (1×10^8^ CFU). (**I**) Schematic model illustrating the regulation of TNF-induced cell death by mTORC1. Data in all graphs are presented as mean ± SD. B shows representative H&E staining from three independent experiments; F shows representative immunofluorescence images from two independent experiments; and E shows representative immunoblots from three independent experiments. Statistical significance was determined using the log-rank (Mantel–Cox) test for survival analyses (A and G), and two-way ANOVA for group comparisons (C, D, H). Significance levels are indicated as follows: ****P < 0.0001; ***P < 0.001; ns, not significant.

We next tested whether this protective mechanism relies on the same ATG9A/FIP200–CHUK/IKKβ-mediated stabilization of later-stage complex I in hepatocytes. In both primary mouse hepatocytes and human HL-7702 liver cells, ATG9A KO or TPCA-1 treatment restored TNF-induced cell death when mTORC1 was inhibited (Figures S7H-S7J). Moreover, both interventions destabilized complex I under these conditions (Figures S7K-S7M).

Beyond inflammatory pathology, TNF-induced cell death contributes to host defense against bacterial infections.^1,32^ Accordingly, hepatocyte-specific Raptor knockout mice showed higher bacterial burdens in the liver and, to a lesser extent, the kidney and spleen, and succumbed more rapidly to *Staphylococcus aureus* infection than WT controls (Figures 7G and 7H). However, additional deletion of ATG9A restored bacterial burdens and mice survival to WT levels (Figures 7G and 7H). Together, these findings demonstrate that mTORC1 inhibition activates an ATG9A/FIP200–CHUK/IKKβ-dependent pathway that stabilizes later-stage complex I to suppress TNF-induced fulminant hepatitis but compromises antibacterial immunity (Figure 7I).

## DISCUSSION

TNF-induced cell death plays a central role in various inflammatory diseases and has long been a focus of therapeutic development.^33–36^ The transition from membrane-bound complex I to cytosolic complex II is required for the propagation of cell death, and elucidating this process may inform pharmacological intervention. We show that mTORC1 suppresses ATG9A/FIP200-dependent accumulation of CHUK/IKKβ, which destabilizes later-stage complex I and facilitates complex II formation. This regulatory mechanism provides insights into TNF signaling modulation, particularly under conditions where mTORC1 activity is diminished.

mTORC1 activity is frequently suppressed or dynamically altered across a wide range of pathological conditions, including cancer, metabolic disorders, inflammatory and neurodegenerative diseases, and infections.^14,37–39^ Among these, TNF-induced cell death plays diverse roles—contributing to tumor suppression, exacerbating inflammatory tissue injury, and enhancing host defense against pathogens. Thus, fluctuations in mTORC1 activity commonly accompany TNF-induced cell death in disease contexts. Our study suggests that the ATG9A/FIP200–CHUK/IKKβ axis, which is inversely regulated by mTORC1 activity, can modulate the progression of TNF-related pathologies. For example, during bacterial infections, mTORC1 is frequently suppressed through various mechanisms.^40–42^ We show that targeting the ATG9A/FIP200–CHUK/IKKβ axis enhances host survival under such conditions. Similarly, in tumors with poor vascularization, mTORC1 activity is often diminished,^43–47^ which may protect cancer cells by impairing TNF-induced cell death. Our findings indicate that inhibiting ATG9A/FIP200–CHUK/IKKβ reactivates TNF-mediated cytotoxicity in these mTORC1-deficient tumor regions.

Unexpectedly, we found that inhibition of mTORC1 leads to stabilization of the TNF-induced later-stage complex I—a regulatory checkpoint not previously described. Although the ATG9A/FIP200–CHUK/IKKβ axis is essential for this stabilization, the precise mechanism remains unclear. We detected no direct interaction between mTORC1 components (Raptor/mTOR) and ATG9A or FIP200, suggesting the involvement of an unidentified bridging substrate. It is also unknown whether this axis functions through a noncanonical role of autophagy proteins or independently of autophagy altogether. Interestingly, we observed that ATG9A expression slightly increased in FIP200-deficient cells, and vice versa (Figure 6F and 6G, input), implying reciprocal regulation. Moreover, since neither ATG9A nor FIP200 are recruited to complex I, the mechanism by which they increase the level of CHUK/IKKβ on complex I (by facilitating CHUK/IKKβ translocation or inhibiting CHUK/IKKβ dissociation) remains to be elucidated. As IKKβ phosphorylates multiple complex I components,^8,26^ identifying the critical substrate(s) and related ubiquitination involved in complex stabilization will be key focuses of future research. Despite these unknowns, our findings reveal a multilayered mechanism regulating the stability of later-stage complex I and suppressing TNF-induced cell death. This has broad therapeutic implications across several disease settings.

## Acknowledgments

We are grateful to Dr. Ding Ai (Tianjin Medical University) for the *Raptor^fl/fl^* mice, Dr. Hui Jiang (National Institute of Biological Sciences, Beijing) for the *Alb-Cre* mice, Dr. Sudan He (Suzhou Institute of Systems Medicine) and Sironax (Beijing) for the *Ripk3^-/-^Caspase8^-/-^* mice, and Dr. Ningning Liu (Institute of Microbiology) for *S. aureus* USA300. We thank Dr. Lilin Du (National Institute of Biological Sciences, Beijing) for providing the beads used in the GFP pulldown assays. We used ChatGPT to correct grammatical errors. This work was supported by the National Natural Science Foundation of China (32321004, 32422023 to Y.A.; 22477095, 22307139 to B.Y.), the National Key R&D Program of China (2022YFA1304500 to Y.A.); and the Beijing Natural Science Foundation Changping Frontier Project (L244054 to Y.A.).

## Author contributions

Y.A. and B.Y. jointly conceptualized and supervised the project. Z.D., B.D., and J.W. performed the majority of the experiments. K.Y. assisted with gene knockout validation and western blot analysis. H.L. contributed to the purification of cytokines for cell-based assays. J.Y. analyzed the CRISPR-Cas9 screen results. H.W. helped with the identification of binding interface of Raptor and Rag. The manuscript was written by Y.A., B.Y., and Z.D., with input from all authors.

## Data and materials availability

All the data supporting the findings of this study are available upon request from.

## MATERIALS AND METHODS

### Cell culture

All mammalian cell lines were maintained at 37 °C in a humidified incubator with 5% CO 2. HeLa, HCT116, MCF-7, MC38, ME180, BMDM, iMEF, and 293T cells were cultured in Dulbecco’s Modified Eagle Medium (DMEM; Gibco, C11995500BT). EMT6 and HL-7702 cells were cultured in RPMI-1640 medium (Gibco, C11875500BT). HeLa, iMEF, MC38, ME180, and HCT116 cells were obtained from the American Type Culture Collection (ATCC). 293T, MCF-7, EMT6, iMEF, and HL-7702 cells were obtained from the Cell Resource Center, Peking Union Medical College (PCRC). All cell lines were routinely tested and confirmed to be free of mycoplasma contamination.

### Animals

All animal experimental protocols were approved by the Animal Research Ethics Committee of Institute of Genetics and Developmental Biology (permit number: AP2025001). *Raptor^fl/fl^* mice were gifted from Prof. Ding Ai of Tianjin Medical University.^1^ *Atg9a^fl/fl^* mice (Strain NO. T022409) were purchased from GemPharmatech (Nanjing, China). *Alb-Cre* mice were gifted from Prof. Hui Jiang of NIBS. *Ripk3^-/-^Caspase8^-/-^* mice were gifted from Dr. Sudan He (SISM) and Sironax (Beijing). All animals used in this study were in the C57BL/6 background. All animals housed in a specific pathogen free facility at constant temperature and humidity under a light cycle of 12 h on/12 h off set from 6 a.m. to 6 p.m.

### Treatment of cells with cytokines and compounds

Various cytokine and compound treatments were applied to different cell lines as indicated. HeLa, HL-7702, ME180, and MCF-7 cells were treated with 50 ng/mL TNF (Cat# 10602-HNAE, Sino Biological) in combination with either 500 nM Smac mimetic (CAS No. 411230-24-5; synthesized in-house), 25 μg/mL cycloheximide (CHX; Cat# 239763-M, Sigma-Aldrich), or 100 ng/mL IFNγ (Cat# 11725-HNAS, Sino Biological). HCT116 cells were treated with 2 ng/mL TNF plus 500 nM Smac mimetic, 50 ng/mL TNF plus 100 ng/mL IFNγ, or 50 ng/mL TNF plus 25 μg/mL CHX. EMT6, BMDM, MC38, and iMEF cells were treated with 50 ng/mL TNF in combination with either 500 nM Smac mimetic or 100 ng/mL IFNγ. For TRAIL-induced apoptosis, HeLa and HCT116 cells were treated with 100 ng/mL TRAIL. Intrinsic apoptosis was induced using 5 μM ABT-737 (Cat# 197333, Sigma-Aldrich) and 2 μM S63845 (Cat# HY-100741, MedChemExpress).

### Cell viability assay

Cell viability was assessed by measuring intracellular ATP levels using the CellTiter-Glo Luminescent Cell Viability Assay (Cat# G7573; Promega), following the manufacturer’s instructions. Luminescence was detected using a PerkinElmer EnSpire microplate reader. Data were analyzed with GraphPad Prism 8.0 (GraphPad Software, San Diego, CA, USA).

### Cell death assay by PI staining

Cells (1 × 10⁵ per well) were seeded in 24-well plates one day prior to treatment. Cells were then subjected to the indicated treatments for the designated durations. For flow cytometry analysis, cells were harvested by trypsinization and resuspended in PBS containing 1 μM propidium iodide (PI; Cat# 537059, Sigma-Aldrich). After a 10-minute incubation at room temperature, samples were analyzed using a flow cytometer (BD FACSAria Fusion), and data were processed with FlowJo software.

### Isolation of Primary Mouse Hepatocytes

Primary mouse hepatocytes were isolated using a two-step collagenase perfusion method. Briefly, 8-12-week mice were anesthetized and perfused via the portal vein with calcium- and magnesium-free HBSS (without phenol red), supplemented with 0.5 nM EDTA and 25 nM HEPES, at 37°C and a flow rate of 5 mL/min. This was followed by perfusion with HBSS containing calcium and magnesium (no phenol red), supplemented with 25 nM HEPES and 1 mg/mL collagenase Type IV (Gibco; 17104019), also at 37°C and 5 mL/min. The digested liver tissue was dissociated in DMEM containing 10% FBS, filtered through a 100 μm mesh, and hepatocytes were purified by centrifugation at 300 rpm for 3 minutes. Viability was assessed by Trypan Blue exclusion, and preparations with >85% viable cells were used. Cells were seeded onto collagen-coated dishes in DMEM supplemented with 10% FBS.

### Western blotting

Cells were lysed either in RIPA buffer (50 mM Tris-HCl pH 7.4, 150 mM NaCl, 1% Triton X-100, 1% sodium deoxycholate, 0.1% SDS, and cOmplete™ EDTA-free Protease Inhibitor Cocktail [Roche, 05892791001]) or directly solubilized in 1 × SDS sample buffer, followed by SDS–polyacrylamide gel electrophoresis (SDS-PAGE). For samples lysed with RIPA buffer, lysates were incubated on ice for 30 minutes and then centrifuged at 15,000 × g for 10 minutes. The supernatants were mixed with 4 × SDS sample buffer and subjected to SDS-PAGE. Proteins were transferred to nitrocellulose membranes (PALL, 66485) using a constant current of 400 mA for 90 minutes. Membranes were blocked with TBST containing 5% non-fat milk for 1 hour at room temperature and incubated overnight at 4°C with primary antibodies diluted in the same buffer. After three 5-minute washes with TBST, membranes were incubated for 1 hour at room temperature with horseradish peroxidase (HRP)-conjugated anti-rabbit or anti-mouse secondary antibodies. After another three washes in TBST, membranes were developed using chemiluminescent substrate (Perkin Elmer, NEL105001EA), and signals were detected using a chemiluminescence imaging system (SINSAGE, MiniChemi 610).

The following antibodies were used for immunoblotting: anti-HSP90 (13171-1-AP; Proteintech), anti-Raptor (2280; Cell Signaling Technology), anti-ATG9A (ab108338; Abcam), anti-FIP200 (17250-1-AP; Proteintech), anti-CHUK (ER30911; HUABIO), anti-IKKβ(2684; Cell Signaling Technology), anti-ATG13 (18258-1-AP; Proteintech), anti-GFP (50430-2-AP; Proteintech), anti-cleaved-Caspase-8 (human) (9496; Cell Signaling Technology), anti-Caspase-8 (4790; Cell Signaling Technology), anti-cleaved-Caspase-8 (mouse) (8592; Cell Signaling Technology), anti-Caspase-3 (9662; Cell Signaling Technology), anti-cleaved-Caspase-3 (9664; Cell Signaling Technology), anti-β-actin (4970; Cell Signaling Technology), anti-GAPDH (HRP-60004; Proteintech), anti-SHARPIN (14626-1-AP; Proteintech), anti-RIPK1 (3493; Cell Signaling Technology), anti-FADD (14906-1-AP; Proteintech), anti-TNFR1 (21574-1-AP; Proteintech), anti-TRADD (15468-1-AP; Proteintech), anti-IκBα (4814; Cell Signaling Technology), anti-p-NF-κB p65 (3033; Cell Signaling Technology), anti-NF-κB p65 (8242; Cell Signaling Technology), anti-p-p38 MAPK (4511; Cell Signaling Technology), anti-p38 MAPK (14064-1-AP; Proteintech), anti-tubulin (10068-1-AP; Proteintech), anti-FLAG (20543-1-AP; Proteintech), anti-Myc (16286-1-AP; Proteintech), HRP-conjugated anti-rabbit IgG (A0545; Sigma-Aldrich), and HRP-conjugated anti-mouse IgG (A9044; Sigma-Aldrich).

### Genome-wide CRISPR/Cas9 screen

The screen was conducted as previously described.^2^ Briefly, the pooled sgRNA library (GeCKO v2; 1000000049, Addgene) was obtained from Addgene. Lentiviral library amplification, packaging, and viral titer determination were performed according to the manufacturer’s instructions. For the genome-wide screen, HeLa-Cas9 cells were generated and seeded in four 15 cm dishes at a density of 5 × 10^6^ cells per dish (total 2 × 10^7^ cells). After 12 hours, cells were infected with the lentiviral sgRNA library at a multiplicity of infection (MOI) of 0.3. At 36 hours post-infection, cells were treated with 10 μg/mL puromycin to eliminate uninfected cells.

Following selection, infected cells were expanded and seeded into twelve 15 cm dishes at 1 × 10^7^ cells per dish. Once the cells reached >90% confluency, six dishes (6 × 10^7^ cells) were treated with TNF combinations, while the remaining six dishes served as untreated controls and were cultured in parallel for the same duration. Surviving cells were collected, expanded, and subjected to two additional rounds of trigger treatment under the same conditions, with the control group passaged concurrently.

After the final treatment, surviving cells from each group were harvested and lysed to extract genomic DNA, which was used as a template for PCR amplification of integrated sgRNA sequences. The resulting PCR products were sequenced using the NovaSeq 6000 platform (Illumina) by ANNOROAD. Sequencing data were processed and analyzed using the MAGeCK algorithm to quantify sgRNA abundance. Differential sgRNA representation between treatment and control groups was calculated, and P-values were assigned to assess positive and negative selection.

### High-throughput drug screening

HeLa cells were seeded in 50 μL of DMEM per well in 384-well plates (Corning, #3707) using an electronic multichannel pipette. Cells were screened against a custom library of 3,097 compounds, including FDA-approved drugs and investigational agents currently in clinical trials. After 12 hours of incubation, 0.5 μL of each compound solution was pin-transferred into assay plates using the Freedom EVO150 automated platform (Tecan), resulting in a final concentration of 10 μM. For combination screening, cells were co-treated with TNF/IFNγ. DMSO served as a negative control, while TNF/IFNγ treatment was used as a positive control on each plate.

After 48 hours of incubation, cell viability was measured using CytoTox-Glo reagent (G9290, Promega; 8 μL/well). Luminescence was recorded using a PerkinElmer EnVision plate reader. The inhibition ratio was calculated as: Inhibition ratio = (sample - negative control mean) / (positive control mean - negative control mean).

### Viral packaging and stable cell line generation

Lentiviral particles were generated by co-transfecting 293T cells with expression plasmids (encoding Raptor, FADD, RIPK1, or CHUK), the packaging plasmid psPAX2, and the envelope plasmid pMD2.G using polyethylenimine (PEI; 24765, Polysciences). After 48 hours, viral supernatants were collected, filtered, and used to transduce target cells in the presence of 5 μg/mL Polybrene (C0351, Beyotime). Twenty-four hours post-transduction, cells were subjected to selection with either 10 μg/mL puromycin (ant-pr-1, InvivoGene) or 10 μg/mL blasticidin (ant-bl-05, InvivoGene) for 7 consecutive days to generate stable cell lines.

### CRISPR–Cas9-mediated gene knockouts

Gene knockout cell lines were generated using CRISPR-Cas9 technology. HeLa, HCT116, MCF-7, EMT6, or iMEF cells were transfected with sgRNA-expressing pX458 plasmids (172221; Addgene) using PEI, following the manufacturer’s instructions. GFP-positive cells were sorted by flow cytometry using a BD FACSAria Fusion cell sorter. Single GFP-positive cells were then seeded into 96-well plates for clonal expansion. Once confluent, clones were transferred to 24-well plates, and successful gene knockouts were confirmed by immunoblotting. A complete list of primers used is provided in Supplementary Table 1.

### Gene silencing by shRNAs

293T cells were co-transfected with the shRNA-containing pLKO.1 lentiviral plasmid, pMD2.G, and psPAX2 using PEI. After 48 hours, the supernatant containing lentiviral particles was collected and used to infect target cells. Infected cells were selected with puromycin 48 hours post-transduction before being used in subsequent experiments. All shRNAs were obtained from the Sigma shRNA clone library and are identified by their corresponding TRC numbers.

### Complex I immunoprecipitation

HeLa and MCF-7 cells were pre-treated as described in the figure legends and subsequently stimulated, or left unstimulated, with one of the following combinations: 200 ng/mL Flag-tagged human TNF (Flag-hTNF) plus 500 nM Smac mimetic and 20 μM z-VAD; 200 ng/mL Flag-hTNF plus 200 ng/mL IFNγ and 20 μM z-VAD; or 200 ng/mL Flag-hTNF plus 25 μg/mL cycloheximide (CHX) and 20 μM z-VAD. Primary mouse hepatocytes were treated with 200 ng/mL mouse TNF (Flag-mTNF), 25 μg/mL CHX, and 20 μM z-VAD for the indicated durations. After treatment, cells were washed twice with ice-cold PBS and lysed in NP-40 lysis buffer (10% glycerol, 0.5% NP-40, 150 mM NaCl, 10 mM Tris-HCl, pH 8.0) supplemented with EDTA-free protease inhibitor cocktail (4693132001; Roche). Lysates were clarified by centrifugation at 20,000 × g for 10 minutes at 4 °C. The resulting supernatants were incubated overnight with Anti-Flag M2 affinity gel (A2220; Sigma-Aldrich). The next day, the beads were washed four times with NP-40 lysis buffer. Immunoprecipitates or whole-cell lysates were then resuspended in 1 × SDS sample buffer and analyzed by SDS-PAGE followed by immunoblotting.

### Complex II immunoprecipitation

HeLa or MCF-7 cells stably expressing FADD-3×Flag or FADD-GFP were pre-treated as indicated in the figure legends for the specified durations. Cells were then washed twice with ice-cold PBS and lysed in NP-40 lysis buffer (10% glycerol, 0.5% NP-40, 150 mM NaCl, 10 mM Tris-HCl, pH 8.0) supplemented with an EDTA-free protease inhibitor cocktail (4693132001; Roche). Lysates were cleared by centrifugation at 20,000 × g for 10 min at 4°C, and the supernatants were incubated overnight at 4°C with anti-GFP magnetic beads. The next day, beads were washed four times with NP-40 lysis buffer. Immunoprecipitates and input lysates were resuspended in 1 × SDS sample buffer, separated by SDS-PAGE, and analyzed by immunoblotting.

### Immunofluorescence

HeLa and MCF-7 cells stably expressing FADD-GFP/BFP or RIPK1-GFP were pre-treated as described in the figure legends for the indicated durations. Cells were fixed with 4% paraformaldehyde for 15 minutes at room temperature, followed by three washes with PBS. Nuclei were stained with Hoechst (C0031; Solarbio) in PBS for 10 minutes, then washed again three times with PBS. Coverslips were stored at 4°C until imaging. Fluorescence images were acquired using a Zeiss LSM 980 confocal microscope equipped with a 63 × oil immersion objective and analyzed using Zeiss Zen software. Manders’ overlap coefficient was calculated with ImageJ. All images shown are representative of at least three independent experiments.

### Exosome collection

HeLa cells were treated with PBS control or TNF/IFNγ for 36 hours. The culture medium was collected and sequentially centrifuged at 1,000 × g for 10 minutes and 10,000 × g for 30 minutes at 4 °C to remove cells and debris. The resulting supernatant was then ultracentrifuged at 140,000 × g for 3 hours at 4 °C to pellet exosomes (Optima L-100XP, Beckman). The pellet was directly resuspended in 1× SDS loading buffer and analyzed by Western blotting. Following medium removal, adherent cells were rinsed once with cold PBS, scraped, and immediately lysed in 1× SDS lysis buffer for Western blot analysis.

### Culture medium collection

HeLa cells were treated with PBS control or TNF/IFNγ for 36 hours. The culture medium was collected and centrifuged at 10,000 × g for 10 minutes at 4 °C. The resulting supernatant was directly mixed with 4 × SDS loading buffer and subjected to Western blot analysis. After medium removal, the adherent cells were washed once with cold PBS, scraped, and immediately lysed in 1 × SDS lysis buffer for Western blotting.

### LC-MS/MS analysis

Proteins were reduced with 10 mM TCEP and alkylated with 40 mM CAA at 45 °C for 5 minutes. Following reduction and alkylation, protein digestion was performed using the SP3 (Single-Pot Solid-Phase-enhanced Sample Preparation) method.^3^ Briefly, 10 μL of Sera-Mag SP3 bead stock (20 mg) was added to each protein sample, followed by addition of acetonitrile to a final concentration of 50% (v/v). Samples were incubated upright at room temperature for 8 minutes. The beads were then washed twice with 200 μL of 70% ethanol, followed by a final rinse with 200 μL of 100% acetonitrile. After discarding the supernatant, proteins were digested in 100 μL of 100 mM TEAB containing trypsin (1:50, w/w) at 37 °C overnight. The resulting tryptic peptides were desalted using StageTips, dried completely in a SpeedVac concentrator, and stored at -20 °C for further analysis.^4^

For MS analysis, peptides were resuspended in 0.1% formic acid (FA) and analyzed using an LTQ Orbitrap Elite mass spectrometer (Thermo Fisher Scientific) coupled online to an Easy-nLC 1000 system in data-dependent acquisition mode. Peptide separation was performed on a 150 μm × 250 mm analytical column packed with 1.9 μm C18 particles using reverse-phase liquid chromatography. The mobile phases were buffer A (0.1% FA) and buffer B (100% acetonitrile, 0.1% FA). Peptides were eluted with a nonlinear gradient of buffer B from 3% to 30% over 90 min. Full MS scans were acquired in the Orbitrap at a resolution of 240,000 (at m/z 400) with a target of 10^6^ ions. The twenty most intense precursor ions were selected for fragmentation by collision-induced dissociation (CID, normalized collision energy: 35) and analyzed in the linear ion trap.

Database searches for all raw MS files were performed using MaxQuant (version 2.3). The *Homo sapiens* reference proteome (20,244 entries) was downloaded from UniProt and used for data searching. Search parameters were set as follows: Type, standard; Multiplicity, 1; Protease, trypsin; Label-free quantification (LFQ), enabled; Minimum score for unmodified peptides, 15. All other parameters were kept at default settings.

### LPS/D-GalN-induced acute liver injury mice models

Male and female C57BL/6 mice (8-10 weeks old) were intraperitoneally injected with lipopolysaccharide (LPS, 10 μg/kg; Sigma, L2630) and D-galactosamine (D-GalN, 300 mg/kg; Macklin, D810561) to induce acute liver injury. Control mice received an equivalent volume of sterile PBS. Four hours post-injection, mice were sacrificed, and liver and blood samples were collected for hematoxylin and eosin (H&E) staining, TUNEL assay, serum AST/ALT measurements, or Western blot analysis, following the manufacturers’ instructions.

### ConA-induced acute liver injury mice models

Male C57BL/6 mice (8-10 weeks old) were used. For survival studies, concanavalin A (ConA; 234-258-2, Sigma) was administered via tail vein injection at 30 mg/kg. For acute liver injury experiments, the dose was reduced to 15 mg/kg. Control mice received an equivalent volume of sterile PBS. Mice were sacrificed 12 hours post-injection, and liver samples were collected for H&E staining and Western blot analysis according to the manufacturer’s instructions.

### *S. aureus* USA300 infections

*S. aureus* USA300 was cultured in LB broth at 37°C. A single colony from an LB agar plate was grown to mid-log phase (OD_600_ = 1), harvested by centrifugation (5,000 × g, 10 min, 4°C), washed with PBS, and resuspended to 2.5 × 10^8^ CFU/mL.

Male and female C57BL/6 mice (8-10 weeks old) were used. For survival studies, mice were injected intravenously via the tail vein with 4 × 10^8^ CFU of *S. aureus* USA300 in 200 μL PBS. For acute infection experiments, the inoculum was reduced to 1 × 10^8^ CFU. Mice were sacrificed 24 hours post-injection, and liver, kidney, and spleen tissues were collected for bacterial burden quantification.

### Bacterial CFU counts

Target organs (e.g., liver, spleen, kidneys) were aseptically harvested, weighed, and homogenized in 4 volumes of PBS using sterile tissue grinders. Homogenates were serially diluted 10-fold in PBS, and 25 μL aliquots were plated onto LB agar. After 24 h incubation at 37 °C, colonies were counted, and bacterial burden was expressed as CFU per gram of tissue (CFU/g).

### Histological analyses

Freshly excised liver tissues were fixed overnight in 4% paraformaldehyde, dehydrated, embedded in paraffin, and sectioned at a thickness of 5 μm. Hematoxylin and eosin (H&E) staining was performed using standard protocols. Images were acquired with a Zeiss Axio Scan.Z1 slide scanner.

### TUNEL assay

Apoptotic cells with DNA fragmentation were detected using the TUNEL assay kit (E-CK-A320; Elabscience) according to the manufacturer’s instructions. Images were captured using a Zeiss LSM 980 confocal microscope.

### ALT and AST Measurements

Serum alanine aminotransferase (ALT) and aspartate aminotransferase (AST) activities were measured to evaluate liver injury using commercial kits. ALT activity was assessed with the Alanine Transaminase Assay Kit (C009-2-1; Nanjing Jiancheng Bioengineering Institute), and AST activity was measured using the Aspartate Transaminase Assay Kit (C010-2-1; Nanjing Jiancheng Bioengineering Institute), following the manufacturers’ protocols.

### Statistical analysis

Statistical analyses were performed using GraphPad Prism 9.0. Data are presented as mean ± SD from three independent experiments. Detailed statistical information is provided in the corresponding figure legends.

---

**Figure S1.**
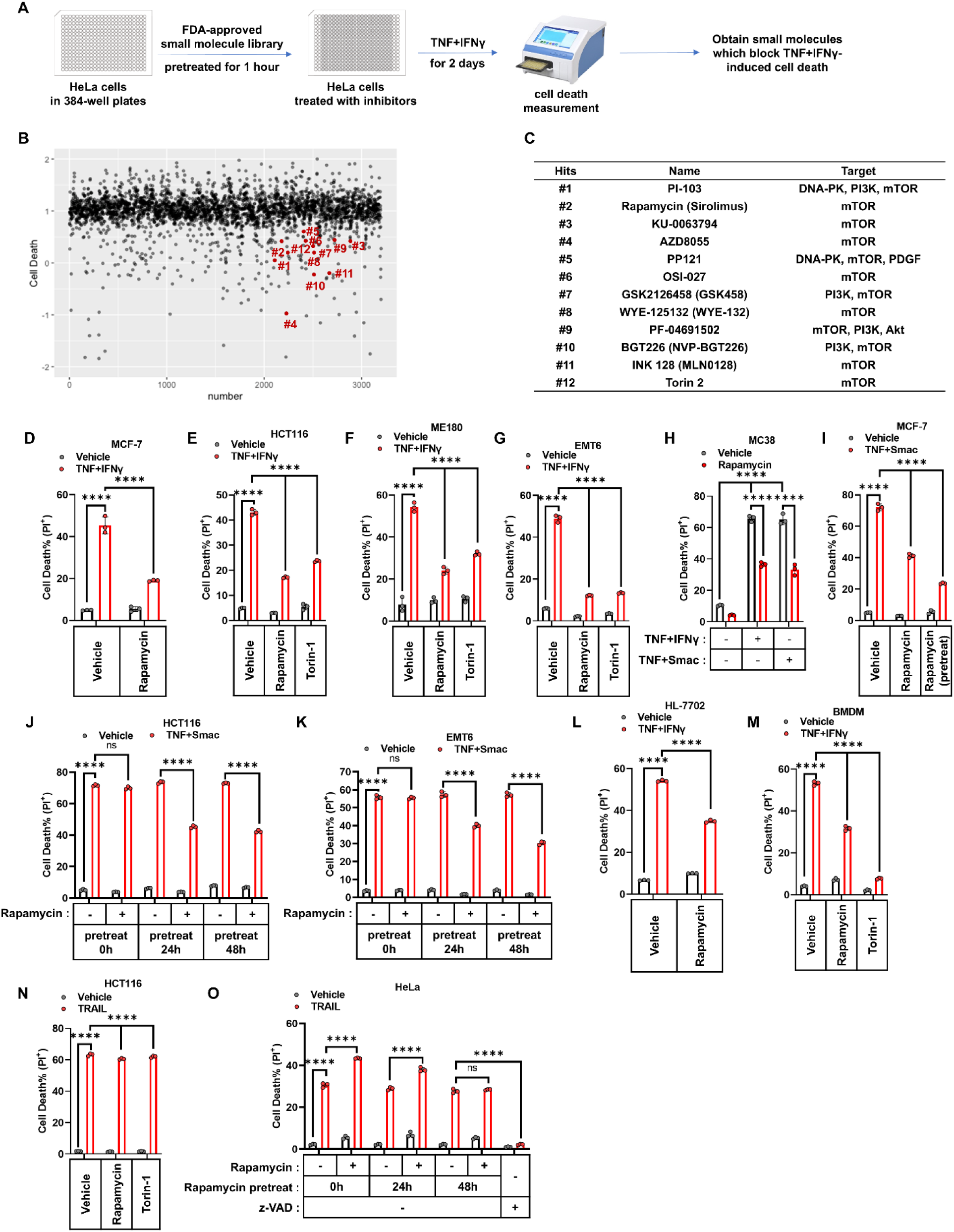
mTORC1 inhibition attenuates TNF-induced cell death. (A) Workflow of high-throughput screening (HTS) in HeLa cells to identify small molecules from an FDA-approved library that block TNF+IFNγ-induced cell death. (B) HeLa cells were treated with compounds from an FDA-approved small molecule library in the presence of TNF+IFNγ. Cell death was quantified by CytoTox-Glo and is shown on the y-axis as normalized inhibition relative to TNF+IFNγ-treated controls. mTOR inhibitors are highlighted as red dots and labeled with corresponding numbers. (C) Names and reported targets of the selected hits highlighted in red in panel B. (D–G) MCF-7 (D), HCT116 (E), ME180 (F), or EMT6 (G) cells were treated with TNF+IFNγ in the presence or absence of the mTOR inhibitors rapamycin (100 nM) or Torin-1 (100 nM). Cell death was assessed by PI staining followed by flow cytometry. (H) MC38 cells were treated with TNF+IFNγ or TNF+Smac in the presence or absence of rapamycin and cell death was assessed. (I) MCF-7 cells were pretreated with rapamycin for 0 h or 24 h followed by TNF+Smac treatment and cell death was assessed. (J, K) HCT116 (J) or EMT6 (K) cells were pretreated with rapamycin for 0 h, 24 h, or 48 h, followed by TNF+Smac treatment and cell death was assessed. (L, M) HL-7702 (L) or BMDM (M) cells were treated with TNF+IFNγ in the presence or absence of rapamycin or Torin-1, and cell death was assessed. (N) HCT116 cells were treated with TRAIL in the presence or absence of rapamycin or Torin-1, and cell death was assessed. (O) HeLa cells were pretreated with rapamycin for 0 h, 24 h, or 48 h, or treated with z-VAD followed by TRAIL, and cell death was assessed. Cell death was assessed by PI staining followed by flow cytometry. The data are presented as mean ± SD, with n = 3 biologically independent samples. Statistical significance in all panels was evaluated using two-way ANOVA: ****P < 0.0001; ns, not significant.

**Figure S2.**
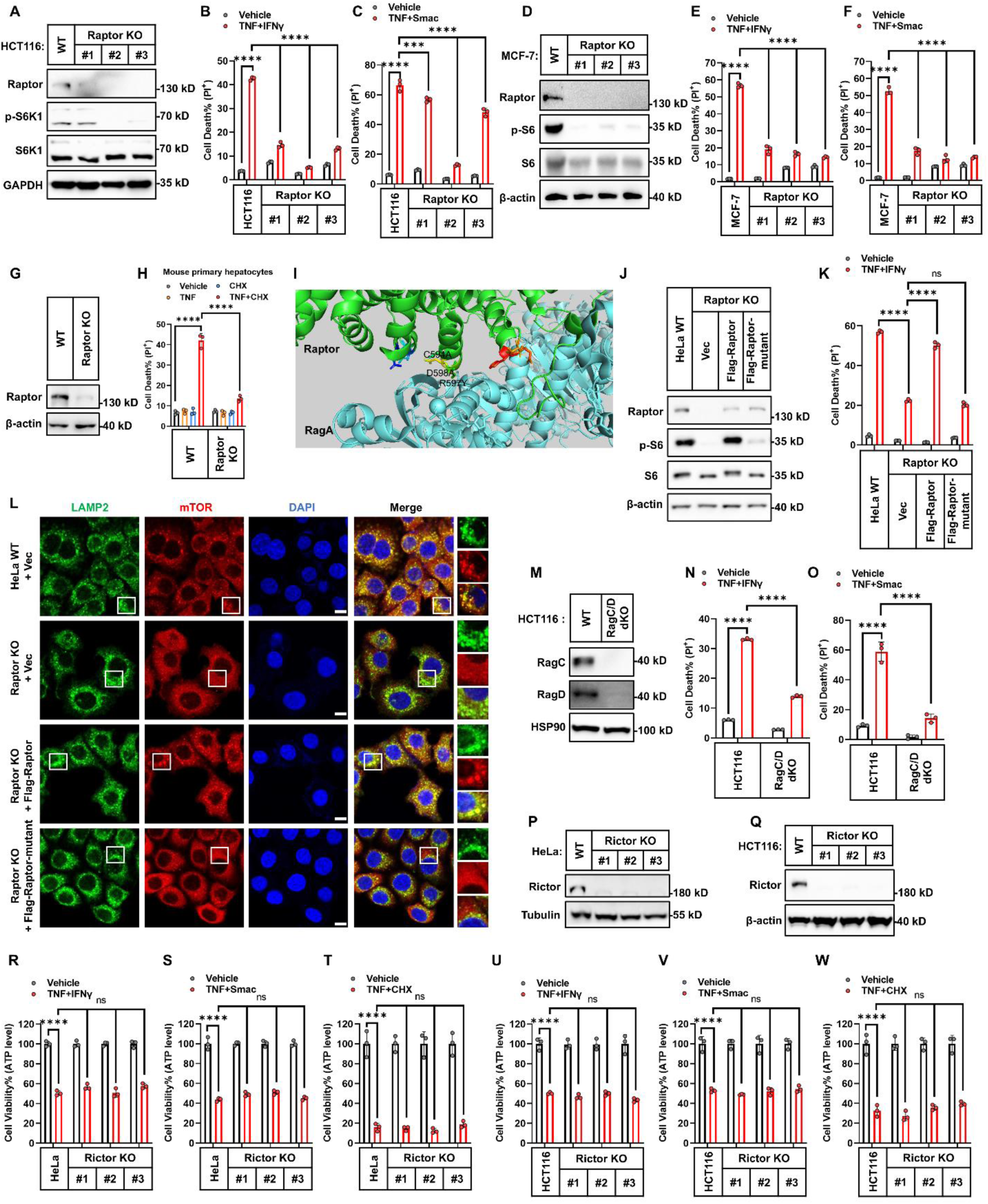
Genetic ablation of mTORC1 impairs TNF-induced cell death across multiple cell types. (A) Raptor KO in HCT116 cells (clones #1, #2, and #3 from different gRNAs) was confirmed by immunoblotting. (B, C) HCT116 cells (parental, Raptor KO clones #1, #2, #3) were treated with TNF+IFN γ (B) or TNF+Smac (C) for 48 h. Cell death was measured. (D) Raptor KO in MCF-7 cells (clones#1, #2 and #3 generated using three different gRNAs) was confirmed by immunoblotting. (E, F). MCF-7 cells (parental, Raptor KO clones #1, #2, #3) were treated with TNF+IFNγ (E) or TNF+Smac (F) for 48 h. Cell death was measured. (G) Raptor KO of mouse primary hepatocytes (WT and Raptor KO) was confirmed by immunoblotting. (H) Mouse primary hepatocytes (WT or Raptor KO) were treated with TNF, CHX, or TNF+CHX for 24 h. Cell death was measured. (I) Structural illustration of the binding interface between Raptor and RagA in the mTOR complex (PDB: 6U62). Critical amino acids involved in the interaction are highlighted. (J) Immunoblot analysis of Raptor and phospho-S6 (p-S6) expression in HeLa parental, Raptor KO, and Raptor KO cells stably reconstituted with either wild-type Flag-Raptor or a Flag-Raptor mutant (C594A/D598A/R597Y). (K) Indicated HeLa cells were treated with TNF+IFNγ for 48 h. Cell death was assessed by PI staining followed by flow cytometry (n = 3). (L) Representative confocal images of the indicated HeLa cells stained for mTOR (green), LAMP1 (lysosomal marker, red), and DAPI (nuclei, blue). Right panels show magnified views of the boxed areas, highlighting mTOR-LAMP1 colocalization. Scale bar: 10 μm. (M) RagC/D dKO in HCT116 cells was confirmed by immunoblotting. (N, O) HCT116 cells (parental and RagC/D dKO) were treated with TNF+IFNγ (N) or TNF+Smac (O) for 48 h. Cell death was measured. (P) Rictor KO in HeLa cells (clones #1, #2, and #3 generated by independent gRNAs) was confirmed by immunoblotting. (Q) Rictor KO in HCT116 cells (clones #1, #2, and #3) was confirmed by immunoblotting. (R-T) HeLa cells (parental and Rictor KO clones) were treated with TNF+IFNγ for 48 h (R), TNF+Smac for 48 h (S), or TNF+CHX for 8 h (T). Cell viability was assessed using an ATP-based luminescence assay (n = 3). (U-W). HCT116 parental and Rictor KO cells were treated with TNF+IFNγ for 48 h (U), TNF+Smac for 48 h (V), or TNF+CHX for 8 h (W). Cell viability was assessed using an ATP-based luminescence assay (n = 3). Cell death in B, C, E, F, H, K, N, O was measured by PI staining followed by flow cytometry. Cell viability in R, S, T, U, V, W was measured using an ATP-based luminescence assay. Data are presented as mean ± SD, with n = 3 biologically independent samples. All immunoblots shown are representative of at least three independent experiments. Statistical significance in all panels was evaluated using two-way ANOVA: ****P < 0.0001; ns, not significant.

**Figure S3.**
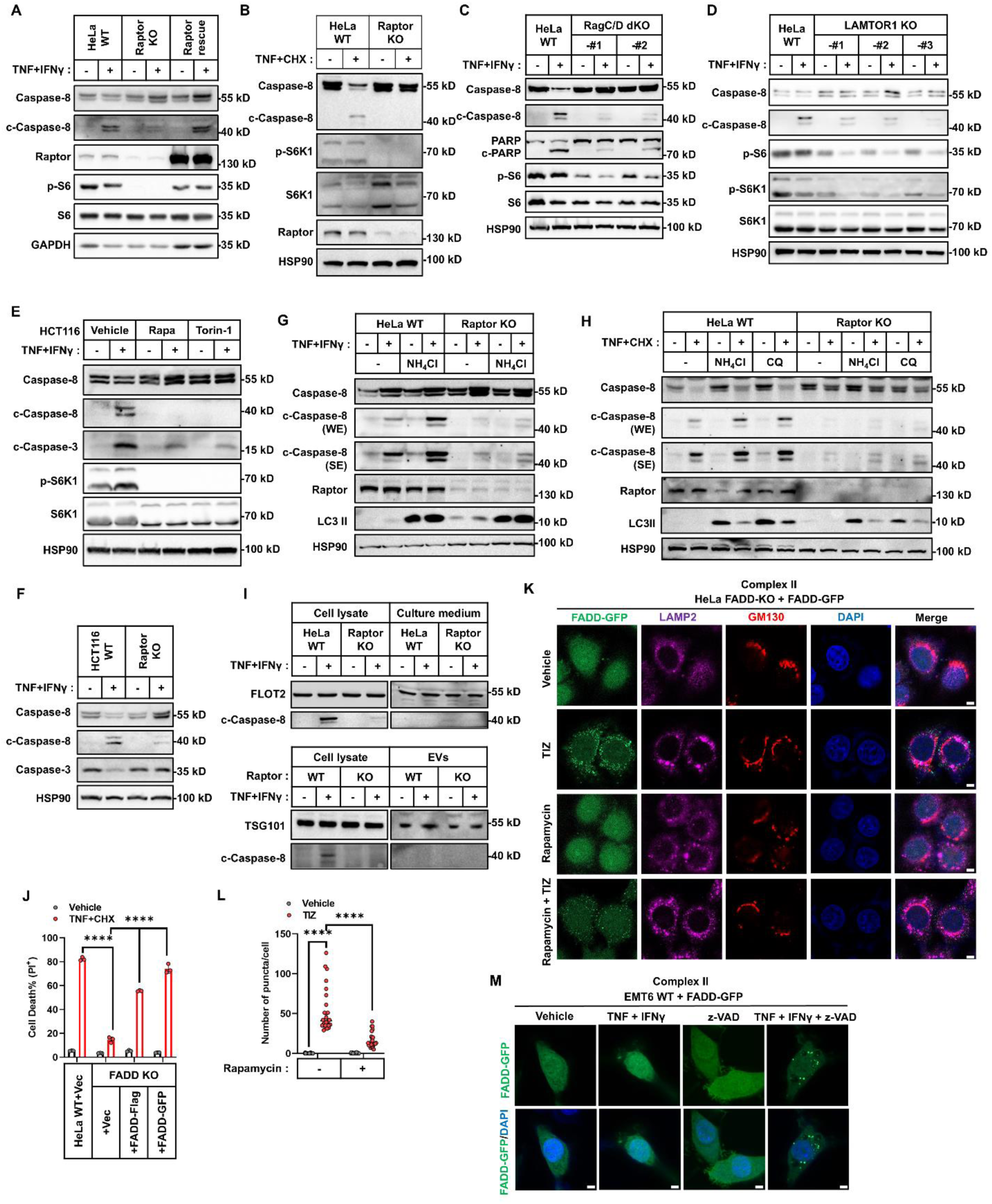
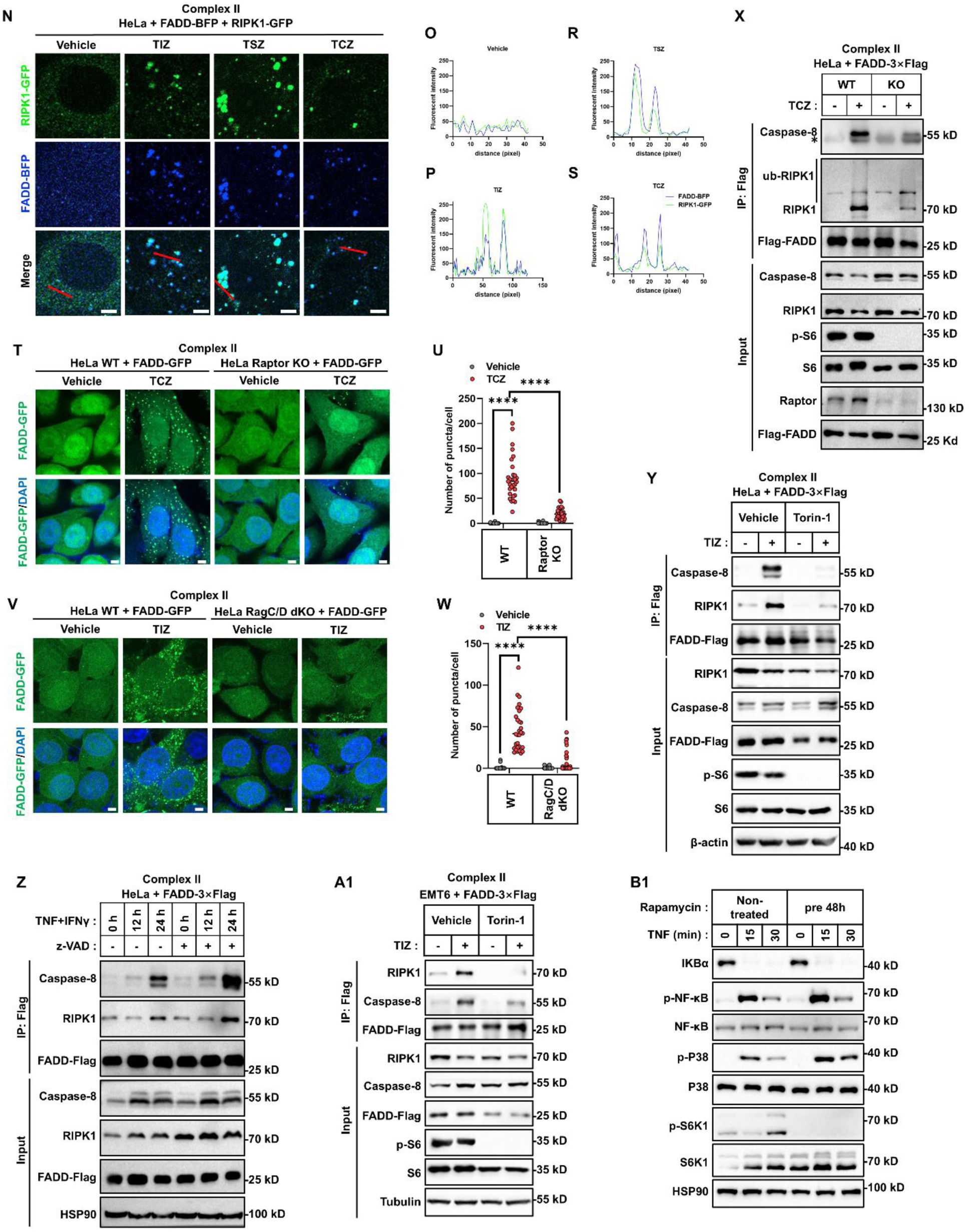
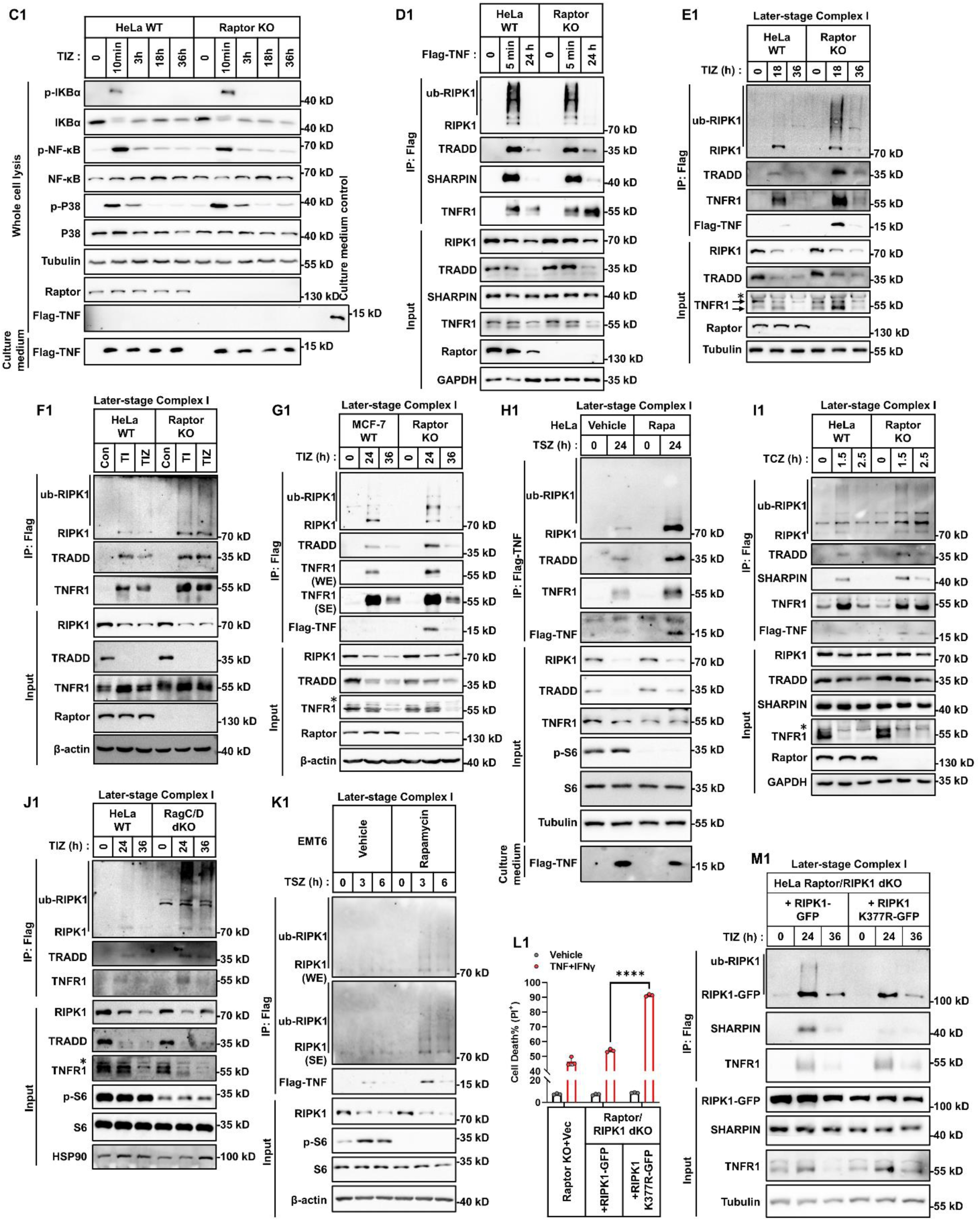
mTORC1 destabilizes membrane-bound later-stage complex I. (A) HeLa cells (parental, Raptor KO, and Raptor KO reconstituted with Raptor) were treated with TNF+IFNγ for 36 h. The expression of the indicated proteins was analyzed by immunoblotting. (B) HeLa cells (parental and Raptor KO) were treated with TNF+CHX for 3 h. The indicated proteins were analyzed by immunoblotting. (C) HeLa cells (parental and RagC/D dKO clones #1 and #2) were treated with TNF+IFN γ for 36 h, and expression of indicated proteins was analyzed by immunoblotting. (D) HeLa cells (parental and LAMTOR1 KO clones #1, #2, and #3) were treated with TNF+IFNγ for 36 h. The expression of indicated proteins was analyzed by immunoblotting. (E) HCT116 cells were treated with TNF+IFNγ in the presence or absence of rapamycin or Torin-1 for 36 h, and indicated proteins was analyzed by immunoblotting. (F) HCT116 cells (parental and Raptor KO) were treated with TNF+IFNγ for 36 h. The levels of indicated proteins were analyzed by immunoblotting. (G) HeLa cells (parental and Raptor KO) were treated with TNF+IFNγ in the presence or absence of NH4Cl for 36 h, and expression of indicated proteins was analyzed by immunoblotting. WE: weak exposure; SE: strong exposure. (H) HeLa cells (parental and Raptor KO) were treated with TNF+CHX in the presence or absence of NH4Cl or CQ for 6 h. The expression of indicated proteins was analyzed via immunoblotting. (I) HeLa cells (parental and Raptor KO) were treated with TNF+IFNγ for 36 h. Cleaved Caspase-8 in cell lysates, culture supernatants, and extracellular vesicles (EVs) was analyzed by immunoblotting. (J) HeLa cells (parental, FADD KO, and FADD KO reconstituted with either FADD-3×Flag or FADD-GFP) were treated with TNF+CHX for 8 h. Cell death was measured by PI staining followed by flow cytometry (n = 3). (K, L) Representative confocal fluorescence images of HeLa FADD KO cells stably expressing FADD-GFP, treated with TIZ for 24 h in the presence or absence of rapamycin. Cells were co-stained with organelle markers (LAMP2 for lysosomes, GM130 for Golgi apparatus) and DAPI (K). Quantification of GFP puncta per cell is shown in (L). Scale bars, 5 μm, n=26 cells. (M) Representative confocal fluorescence images of EMT6 cells stably expressing FADD-GFP treated with TNF+IFNγ in the presence or absence of z-VAD for 24 h. Nuclei were stained with DAPI. Scale bars, 5 μm. (N) Representative confocal fluorescence images of HeLa cells stably expressing FADD-BFP and RIPK1-GFP, treated with TIZ for 24 h, TNF+Smac+z-VAD (TSZ) for 24 h, or TNF+CHX+z-VAD (TCZ) for 2.5 h. Nuclei were stained with DAPI. Scale bars, 5 μm. (O-S) Fluorescence intensity histograms showing grayscale value distribution for the images in (N). (T, U) Representative confocal images of HeLa cells and Raptor KO cells stably expressing FADD-GFP treated with TCZ for 2.5 h. Cytosolic complex II formation was visualized as GFP-positive puncta; nuclei were counterstained with DAPI (T). Quantification of puncta per cell is shown in (U). Scale bar, 5 μm, n=33 cells. (V, W) Representative confocal images of HeLa and RagC/D dKO cells stably expressing FADD-GFP treated with TIZ for 24 h. Complex II puncta and DAPI-stained nuclei are shown (V). Quantification in (W). Scale bar, 5 μm, n=32 cells. (X) HeLa WT and Raptor KO cells stably expressing FADD-3×Flag were treated with TCZ for 2.5 h. Cytosolic complex II was immunoprecipitated using anti-Flag beads, and associated proteins were analyzed by immunoblotting. Asterisk indicates non-specific bands. (Y) HeLa-FADD-3×Flag cells were treated with TIZ with or without Torin-1 for 24 h. Cytosolic complex II was immunoprecipitated using anti-Flag beads, and associated proteins were analyzed via immunoblotting. (Z) HeLa-FADD-3×Flag cells were treated with TNF+IFNγ with or without z-VAD for the indicated times. Cytosolic complex II was immunoprecipitated and analyzed by immunoblotting. (A1) EMT6-FADD-3×Flag cells were treated with TIZ in the presence or absence of Torin-1 for 24 h. Cytosolic complex II was immunoprecipitated and associated proteins were analyzed. (B1) HeLa cells were treated with TNF for the indicated times following pre-treatment with rapamycin (100 nM) for 48 h. Activation of the NF-κB and MAPK signaling pathways was assessed by immunoblotting. (C1) HeLa parental or Raptor KO cells were treated with Flag-TNF for the indicated times. Activation of the NF-κB and MAPK pathways, along with detection of Flag-TNF in cell lysates and culture supernatants, was analyzed by immunoblotting. (D1) HeLa parental or Raptor KO cells were treated with Flag-TNF for the indicated times. Membrane-bound complex I was immunoprecipitated using anti-Flag beads, and associated proteins were examined via immunoblotting. (E1) HeLa cells (parental or Raptor KO) were treated with Flag-TNF+IFNγ+z-VAD (TIZ) for the indicated times. Membrane-bound complex I was immunoprecipitated using anti-Flag antibody, and representative proteins were analyzed by immunoblotting. Asterisk indicates non-specific bands. (F1) HeLa cells (parental or Raptor KO) were treated with Flag-TNF+IFNγ+z-VAD (TIZ) for 24 h. Membrane-bound complex I was analyzed. (G1) MCF-7 cells (parental or Raptor KO) were treated with Flag-TNF+IFNγ+z-VAD (TIZ) for the indicated times, and complex I was analyzed. WE: weak exposure; SE: strong exposure. (H1) HeLa cells were treated with Flag-TNF+Smac+z-VAD (TSZ) for 24 h in the presence or absence of rapamycin (100 nM). Complex I was analyzed. (I1) HeLa cells (parental or Raptor KO) were treated with Flag-TNF+CHX+z-VAD (TCZ) for the indicated times. Membrane-bound complex I was analyzed. Asterisk in immunoblot image indicates non-specific bands. (J1) HeLa cells (parental or RagC/D dKO) were treated with Flag-TNF+IFNγ+z-VAD (TIZ) for the indicated times., and complex I was analyzed. Asterisk indicates non-specific bands. (K1) EMT6 cells were treated with TSZ for the indicated times. Membrane-bound complex I was determined. WE: weak exposure; SE: strong exposure. (L1) Raptor/RIPK1 dKO HeLa cells stably expressing RIPK1-GFP or the ubiquitination-deficient mutant RIPK1-K377R-GFP were treated with TNF+IFNγ for 48 h. Cell death was assessed by PI staining followed by flow cytometry (n = 3). (M1) The same cell lines as in (L1) were treated with Flag-TNF+IFNγ+z-VAD (TIZ) for the indicated times. Membrane-bound complex I was analyzed. Vehicle was used as control for specific stimuli. Data are presented as mean ± SD, with n = 3 biologically independent samples. All representative immunoblots are from two to three independent experiments. Statistical significance was evaluated using the Kruskal–Wallis test (L, U, W) or two-way ANOVA (J, L1): ****P < 0.0001; ns, not significant.

**Figure S4.**
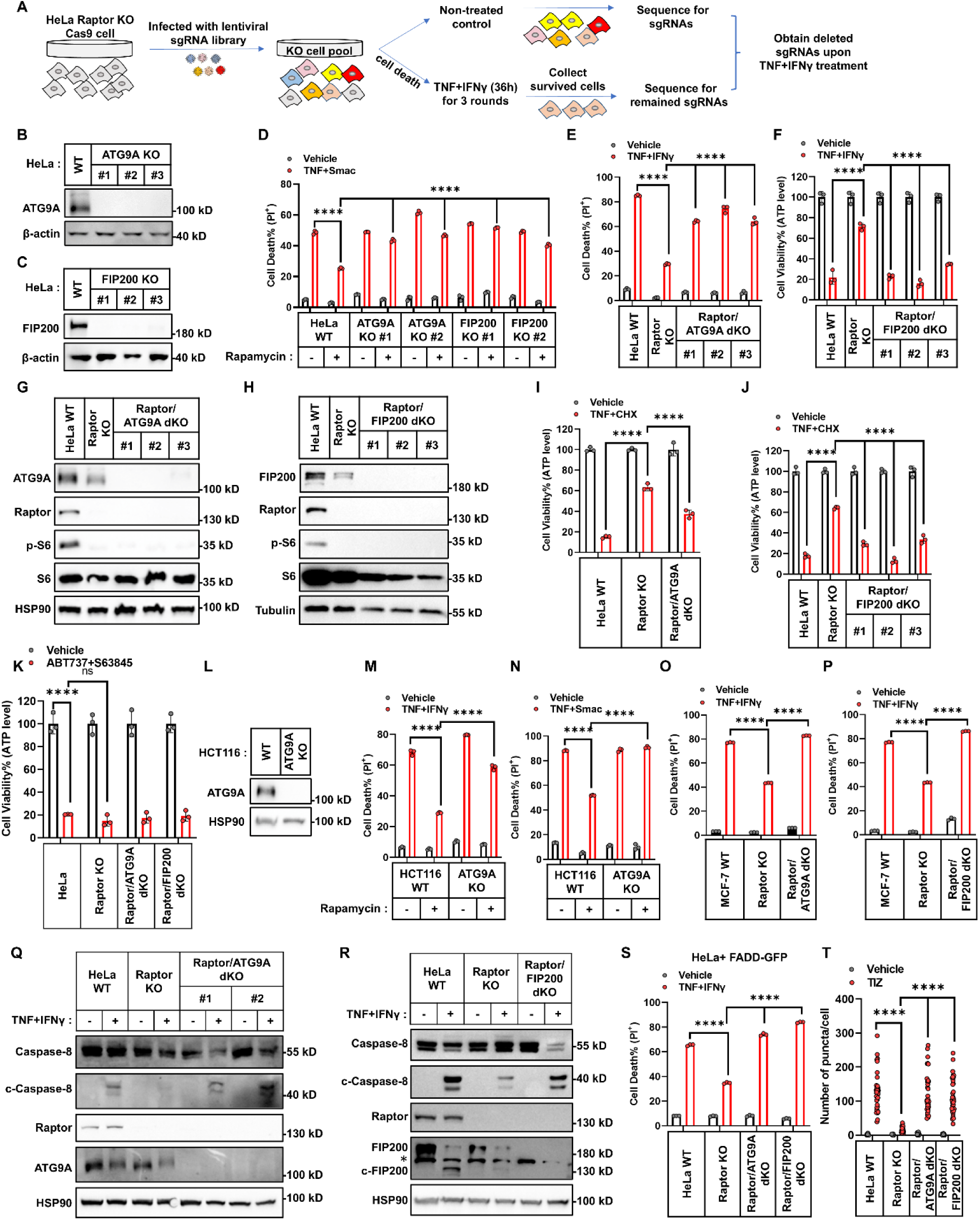
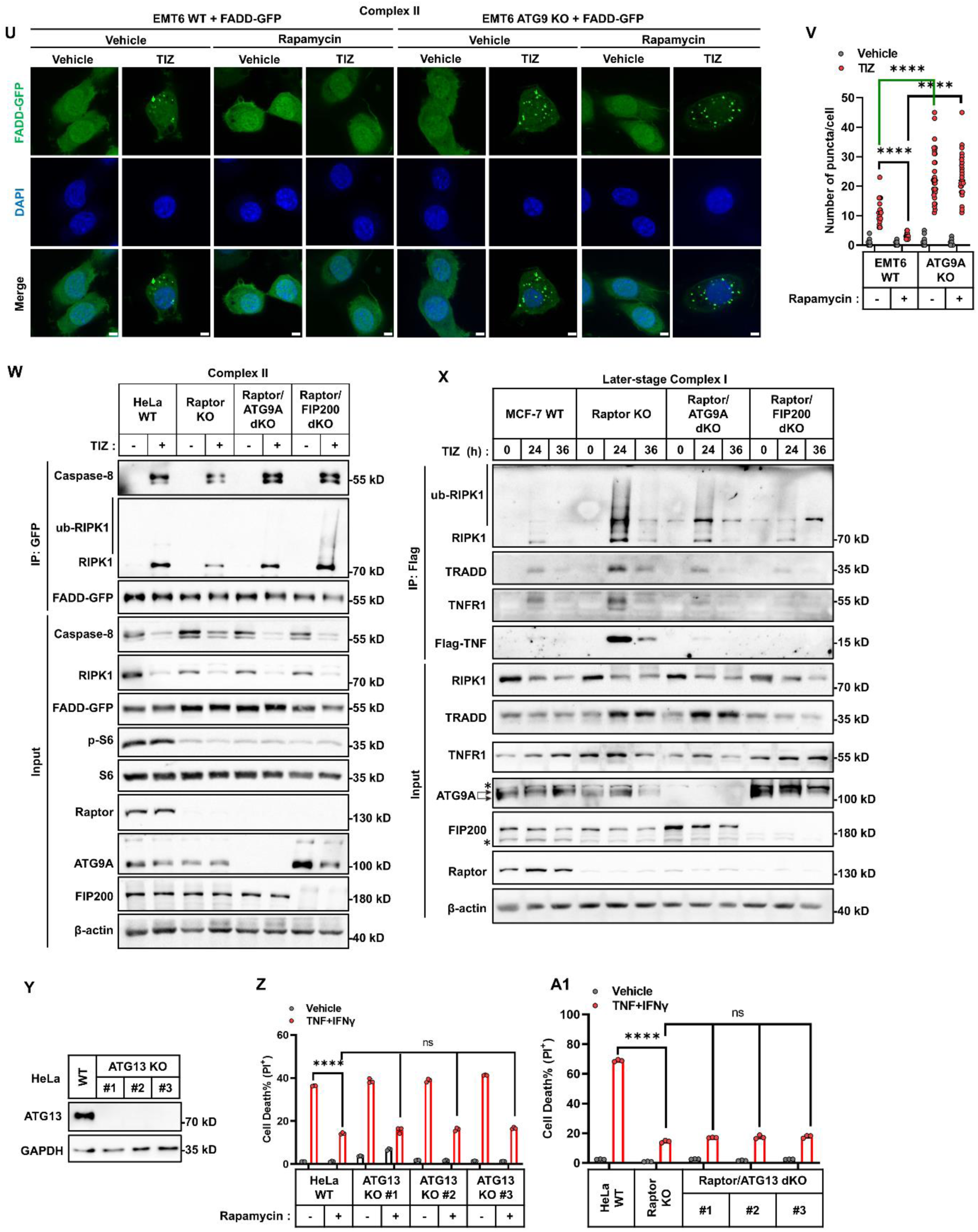
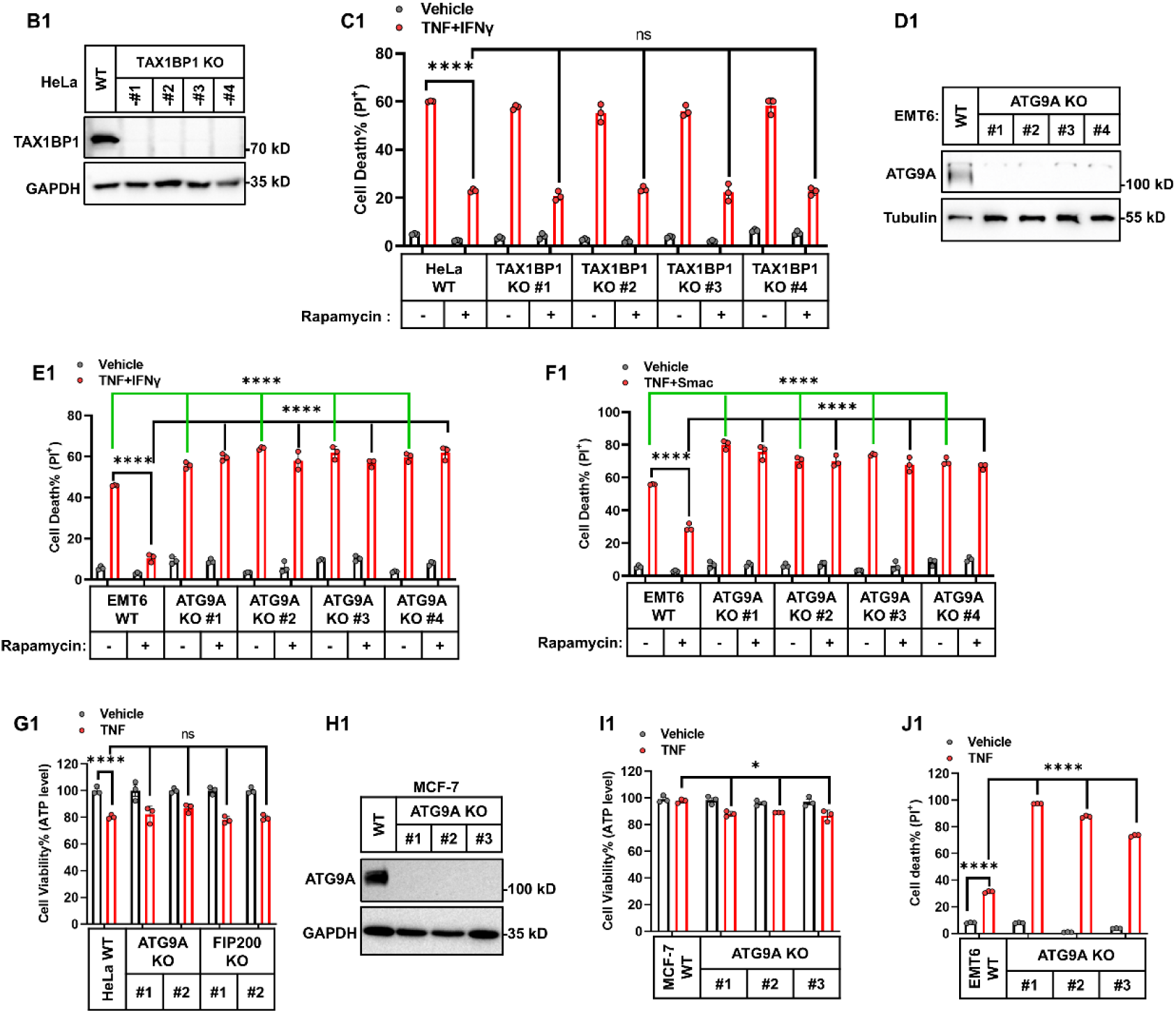
ATG9A and FIP200 are required to suppress complex II formation and stabilize later-stage complex I upon mTORC1 inhibition. (A) Schematic diagram of the CRISPR-Cas9 dropout screening strategy used to identify regulators that inhibit TNF+IFNγ-induced cell death in Raptor KO cells. (B, C) Knockout of ATG9A (B) or FIP200 (C) in HeLa cells (clones #1, #2, and #3 from three different gRNAs) was confirmed by immunoblotting. (D) HeLa ATG9A KO and FIP200 KO cells (clones #1 and #2) were treated with TNF+Smac in the presence or absence of rapamycin (100 nM). Cell death was measured by PI staining. (E) HeLa cells (parental, Raptor KO, and Raptor/ATG9A dKO clones #1, #2, #3) were treated with TNF+IFNγ for 48h. Cell death was measured. (F) HeLa cells (parental, Raptor KO, Raptor/FIP200 dKO clones #1, #2, #3) were treated with TNF+IFNγ for 48h. Cell viability was measured by ATP-based luminescence assay (n = 3). (G, H) Knockout of ATG9A (G) or FIP200 (H) in HeLa Raptor KO cells (clones #1, #2, and #3) was confirmed by immunoblotting. (I, J) HeLa cells (parental, Raptor KO, and Raptor/ATG9A dKO) (I) or (parental, Raptor KO, and Raptor/FIP200 dKO clones) (J) were treated with TNF+CHX for 8 h. Cell viability was measured using an ATP-based luminescence assay (n = 3). (K) HeLa cells (parental, Raptor KO, Raptor/ATG9A dKO, or Raptor/FIP200 dKO clones) were treated with ABT737+S63845 for 8 h. Cell viability was measured (n = 3). (L) Knockout of ATG9A in HCT116 cells was confirmed by immunoblotting. (M, N) HCT116 cells (parental or ATG9A KO) were treated with TNF+IFNγ (M) or TNF+Smac (N) for 48 h. Cell death was measured by PI staining. (O, P) MCF-7 cells (parental, Raptor KO, and Raptor/ATG9A dKO) (O) or (parental, Raptor KO, and Raptor/FIP200 dKO) (P) were treated with TNF+IFNγ for 48 h. Cell death was measured (n = 3). (Q) HeLa cells (parental, Raptor KO, and Raptor/ATG9A dKO clones #1 and #2) were treated with TNF+IFNγ for 36 h. The level of cleaved and full-length Caspase-8 was analyzed by immunoblotting. (R) HeLa cells (parental, Raptor KO, and Raptor/FIP200 dKO) were treated with TNF+IFN γ for 36 h. The level of cleaved and full-length Caspase-8 was analyzed. Asterisk indicates non-specific bands. (S) HeLa cells (parental, Raptor KO, Raptor/ATG9A dKO, and Raptor/FIP200 dKO), stably expressing FADD-GFP, were treated with TI for 48 h. Cell death was measured by PI staining followed by flow cytometry (n = 3). (T) Quantification of GFP-positive puncta per cell corresponding to the confocal images in Figure 4E, n=32 cells. (U, V) EMT6 cells (parental or ATG9A KO clone) stably expressing FADD-GFP were treated with TIZ in the presence or absence of rapamycin for 24 h. Cytosolic complex II formation was visualized by confocal microscopy as GFP-positive puncta. Nuclei were counterstained with DAPI (U). GFP puncta per cell was quantified (V). Scale bar, 5 μm, n=33 cells. (W) HeLa cells (parental, Raptor KO, Raptor/ATG9A dKO, Raptor/FIP200 dKO) stably expressing FADD-GFP were treated with TIZ for 24 h. Cytosolic complex II was immunoprecipitated using anti-GFP antibody, and co-precipitated proteins were analyzed by immunoblotting. (X) MCF-7 cells (parental, Raptor KO, Raptor/ATG9A dKO, Raptor/FIP200 dKO) were treated with Flag-TNF+IFNγ+z-VAD (TIZ) for the indicated times. Membrane-bound complex I was immunoprecipitated using anti-Flag antibody, and co-precipitated proteins were analyzed by immunoblotting. Asterisk indicates non-specific bands. (Y) ATG13 KO in HeLa cells (clones #1, #2, and #3 generated using three different gRNAs) was confirmed by immunoblotting. (Z) HeLa cells (parental, ATG13 KO clones #1, #2, #3) were treated with TNF+IFN γ in the presence or absence of rapamycin (100 nM) for 48 h. Cell death was measured by PI staining followed by flow cytometry (n = 3). (A1) HeLa cells (parental, Raptor KO, and Raptor/ATG13 double KO clones #1, #2, #3) were treated with TNF+IFNγ for 48 h. Cell death was measured (n = 3). (B1) TAX1BP1 KO in HeLa cells (clones #1, #2, #3, and #4 generated using four different gRNAs) was confirmed by immunoblotting. (C1) HeLa cells (parental, TAX1BP1 KO clones #1, #2, #3, #4) were treated with TNF+IFN γ in the presence or absence of rapamycin (100 nM) for 48 h. Cell death was measured as above (n = 3). (D1) ATG9A KO in EMT6 cells (clones #1, #2, #3, and #4 generated using four different gRNAs) was confirmed by immunoblotting. (E1, F1) EMT6 cells (parental, ATG9A KO clones #1, #2, #3, #4) were treated with TNF+IFN γ (E1) for 48 h or TNF+Smac (F1) for 24 h in the presence or absence of rapamycin. Cell death was measured by PI staining followed by flow cytometry (n=3). (G1) HeLa cells (parental, ATG9A KO clones #1, #2, FIP200 KO clones #1, #2) were treated with TNF for 48 h. Cell viability was measured via ATP-based luminescence assay (n = 3). (H1) ATG9A KO in MCF-7 cells (clones#1, #2, #3 from three different gRNAs) was confirmed by immunoblotting. (I1) MCF-7 cells (parental, ATG9A KO clones #1, #2, #3) were treated with TNF for 48 h. Cell viability was measured (n=3). (J1) The mouse cell line EMT6 cells (parental, ATG9A KO clones #1, #2, #3) were treated with TNF for 48 h. Cell death was measured by PI staining (n=3). The data are presented as mean ± SD, with n = 3 biologically independent samples. Panel E shows representative immunofluorescence from two independent experiments; all immunoblots shown are representative of at least two independent experiments. Statistical significance was evaluated using the Kruskal–Wallis test (T, V) or two-way ANOVA: ****P < 0.0001; ns, not significant.

**Figure S5.**
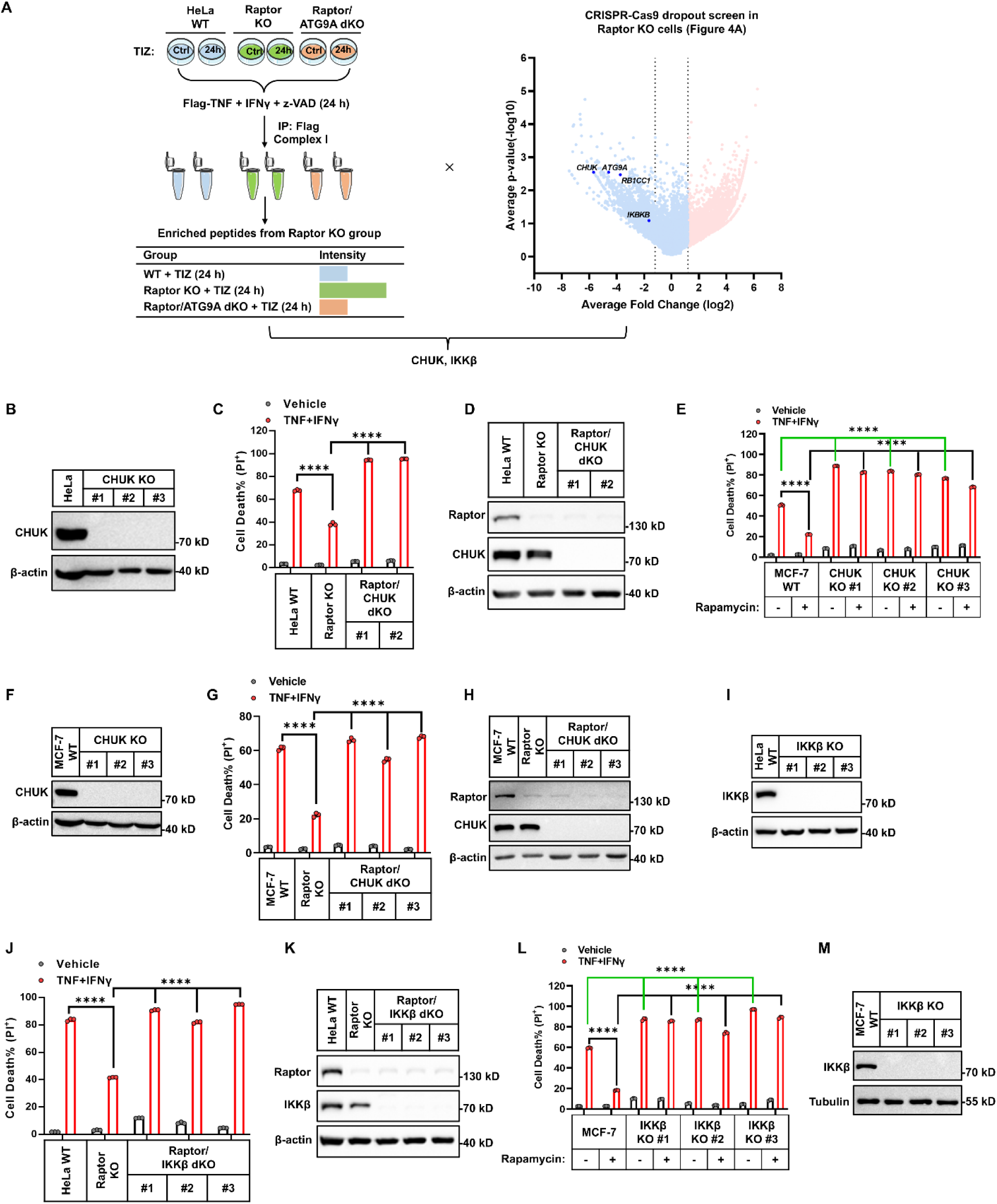
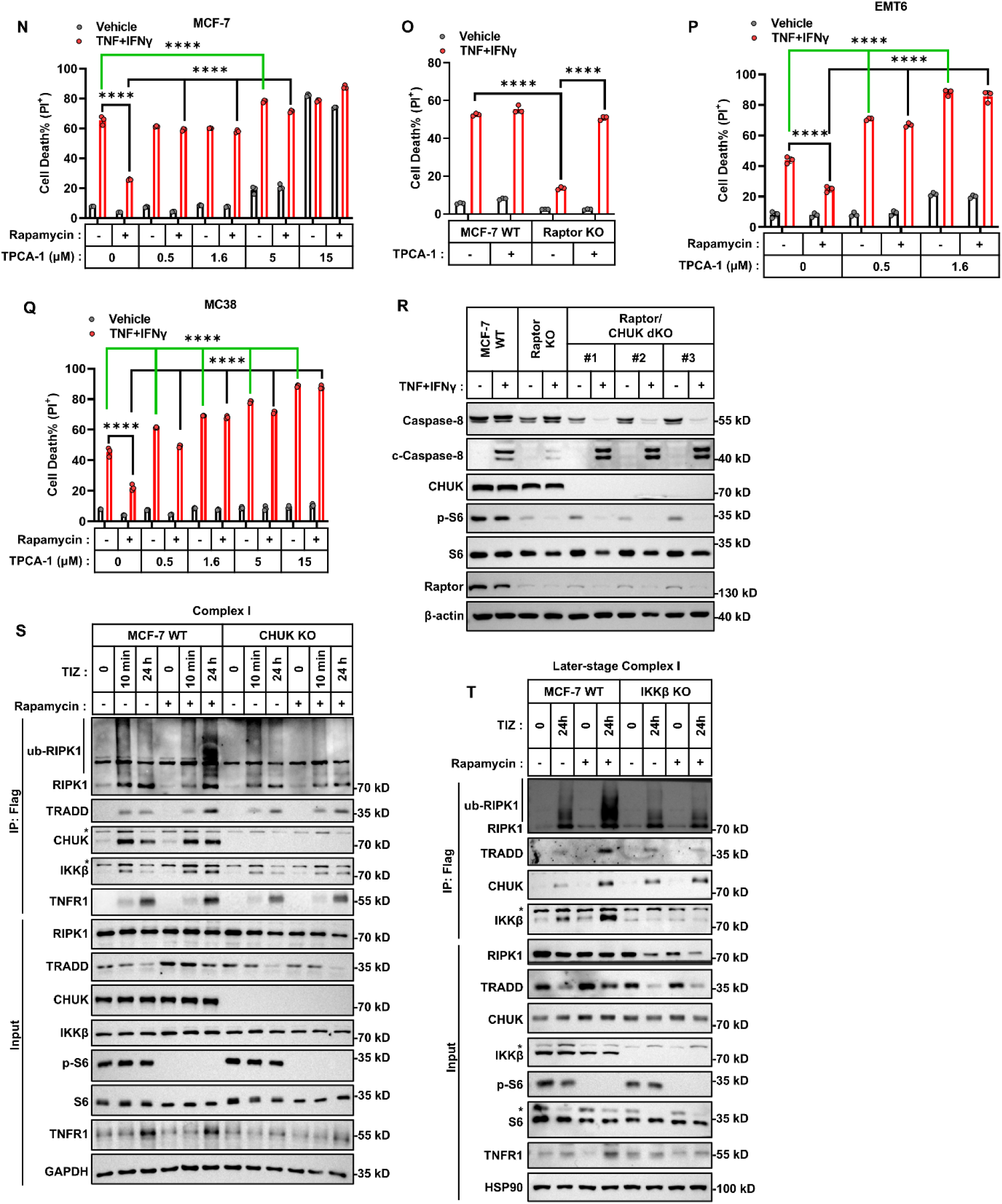
CHUK/IKKβ mediates the protective effect of mTORC1 inhibition on TNF-induced cell death. (A) Schematic overview of the LC/MS workflow is shown on the left: HeLa parental, Raptor KO, and Raptor/ATG9A dKO cells were treated with TNF+IFNγ for 0 or 24 h, followed by immunoprecipitation of complex I and mass spectrometry analysis. Proteins enriched in Raptor KO cells were considered candidates. On the right, the CRISPR-Cas9 dropout screen from Figure 4A was used to identify overlapping candidates from both LC/MS and genetic screening datasets. (B) CHUK KO in HeLa cells (clones #1, #2, #3 from three different gRNAs) was confirmed by immunoblotting. (C) HeLa cells (parental, Raptor KO, Raptor/CHUK dKO clones #1, #2) were treated with TNF+IFNγ for 48 h. Cell death was measured by PI staining followed by flow cytometry (n=3). (D) CHUK KO in HeLa Raptor KO cells (clones #1, #2) was confirmed. (E) MCF-7 cells (parental, CHUK KO clones #1, #2, #3) were treated with TNF+IFNγ in the presence or absence of rapamycin for 48 h. Cell death was measured by PI staining. (F) CHUK KO in MCF-7 cells was confirmed via immunoblotting. (G) MCF-7 cells (parental, Raptor KO, Raptor/CHUK dKO clones #1, #2, #3) were treated with TNF+IFNγ for 48 h. Cell death was measured (n=3). (H) CHUK KO in MCF-7 Raptor KO cells was confirmed. (I) IKKβ KO in HeLa cells (clones #1, #2, #3 from three different gRNAs) was confirmed by immunoblotting. (J) HeLa cells (parental, Raptor KO, Raptor/IKKβ dKO clones #1, #2, #3) were treated with TNF+IFNγ for 48 h. Cell death was measured (n=3). (K) IKKβ KO in HeLa Raptor KO cells was confirmed via immunoblotting. (L) MCF-7 cells (parental, IKKβ KO clones#1, #2, #3) were treated with TNF+IFNγ in the presence or absence of rapamycin for 48 h, and cell death was measured (n=3). (M) IKKβ KO in MCF-7 cells was confirmed by immunoblotting. (N) MCF-7 cells were treated with TNF+IFNγ in the presence or absence of rapamycin with different concentrations of TPCA-1 for 48 h, and cell death was measured. (O) MCF-7 cells (parental, Raptor KO) were treated with TNF+IFNγ in the presence or absence of TPCA-1 (0.5μM) for 48 h (n = 3). Cell death was measured. (P, Q) EMT6 (P) or MC38 (Q) cells were treated with TNF+IFNγ in the presence or absence of rapamycin with different concentrations of TPCA-1 for 48 h. Cell death was measured (n=3). (R) Immunoblot analysis of Caspase-8 and cleaved Caspase-8 in MCF-7 WT, Raptor KO, and Raptor/CHUK dKO cells treated with TNF+IFNγ. (S) MCF-7 cells were treated with Flag-TNF+IFNγ+z-VAD (TIZ) in the presence or absence of rapamycin for 10 minutes or 24 h. Membrane-bound complex I was analyzed by immunoprecipitation and immunoblotting. (T) MCF-7 cells (parental or IKKβ KO) were treated with Flag-TNF+IFNγ+z-VAD (TIZ) with or without rapamycin for 24 h. Complex I was determined. The data are showed as mean ± SD, with n = 3 biologically independent samples. All representative immunoblots were obtained from at least two independent experiments. Statistical significance in all panels was evaluated using two-way ANOVA: ****P < 0.0001; ns, not significant.

**Figure S6.**
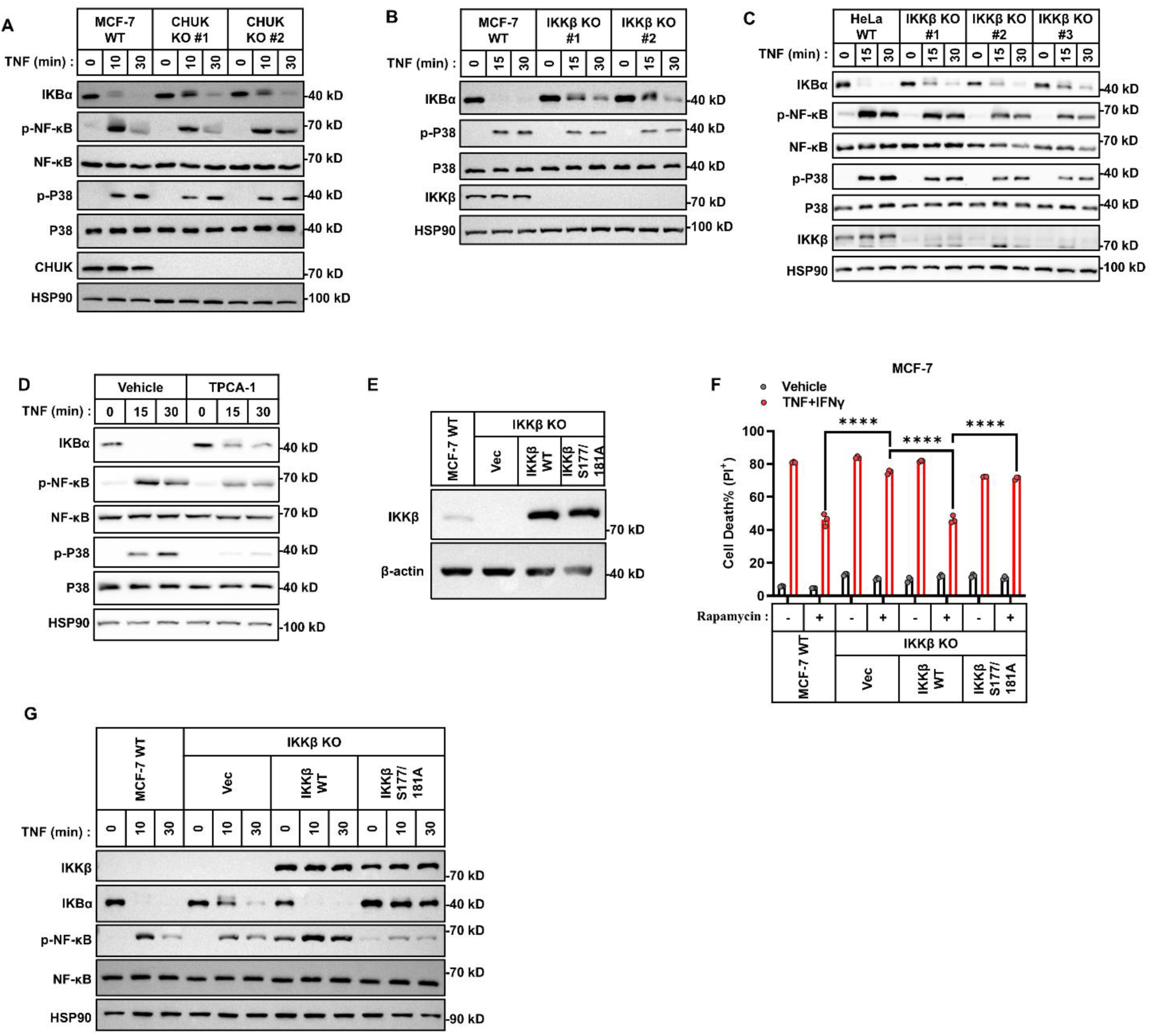
CHUK/IKKβ stabilizes later-stage complex I and suppresses Caspase-8 activation independent of NF-κB signaling. (A) Immunoblotting analysis of NF-κB activation in MCF-7 WT and CHUK KO cells treated with TNF for 0, 10, or 30 minutes. (B, C) Immunoblotting of NF-κB activation in WT and IKKβ KO MCF-7 (B) or HeLa (C) cells treated with TNF for 0, 15, or 30 minutes. (D) HeLa cells were pretreated with or without TPCA-1 followed by TNF treatment for the indicated times. NF-κB and MAPK activation was assessed by immunoblotting. (E) Immunoblotting analysis of NF-κB activation in MCF-7 WT, IKKβ KO cells were stably reconstituted with either an empty vector, WT IKKβ, or kinase-deficient IKKβ mutants (S1177A and S181A). (F) The indicated MCF-7 cells were treated with TNF+IFNγ for 48 h. Cell death was assessed by PI staining and flow cytometry (n = 3) (G) Immunoblotting analysis of NF-κB activation in MCF-7 WT, IKKβ KO cells were stably reconstituted with either an empty vector, WT IKKβ, or kinase-deficient IKKβ mutants (S1177A and S181A), cells treated with TNF for 0, 10, or 30 minutes. The data are showed as mean ± SD, with n = 3 biologically independent samples. All representative immunoblots are from two to three independent experiments. Statistical significance in all panels was evaluated using two-way ANOVA: ****P < 0.0001; ns, not significant.

**Figure S7.**
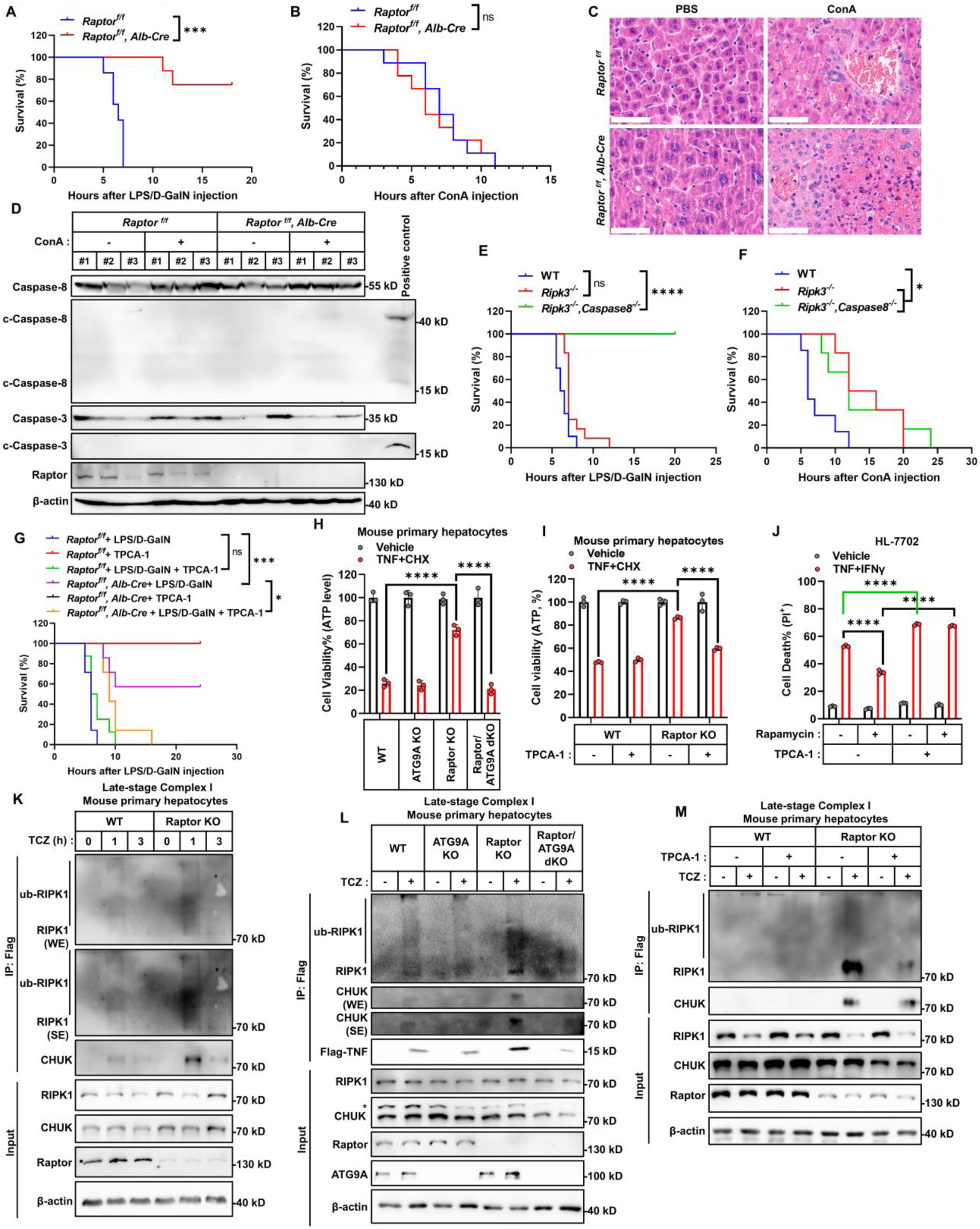
Hepatocyte-specific Raptor deletion protects against TNF-mediated liver injury via ATG9A-dependent stabilization of later-stage complex I but increases susceptibility to bacterial sepsis. (A) Cumulative survival curves of female *Raptor^fl/fl^* and *Raptor^fl/fl^ Alb-Cre* C57BL/6J mice (8 weeks old) intraperitoneally injected with LPS/D-GalN (n = 10). (B) Cumulative survival curves of male *Raptor^fl/fl^* and *Raptor^fl/fl^ Alb-Cre* mice (8 weeks old) intraperitoneally injected with ConA (n = 10). (C) Representative H&E staining of liver sections from *Raptor^fl/fl^* and *Raptor^fl/fl^ Alb-Cre* mice 12 h after ConA injection (n = 6). Scale bar: 50 μm. (D) Immunoblot analysis of cleaved Caspase-8 and Caspase-3 in liver lysates of indicated mice (n = 3). (E, F) Cumulative survival curves of WT*, Ripk3^-/-^*and *Ripk3^-/-^ Caspase8^-/-^* male mice (8 weeks old) intraperitoneally injected with LPS/D-GalN (E) or ConA (F) (n = 10). (H) Cumulative survival curves of male *Raptor^fl/fl^* and *Raptor^fl/fl^ Alb-Cre* C57BL/6J mice (8 week) intraperitoneally injected with LPS/D-GalN, with or without TPCA-1 (10 mg/kg) (n = 10). (I) Mouse primary hepatocytes (WT, ATG9A KO, Raptor KO, and Raptor/ATG9A dKO) were treated with TNF+CHX for 24 h, and cell viability was assessed using ATP-based luminescence assay (n = 3). (J) Mouse primary hepatocytes (WT and *Raptor* KO) were treated with TNF+CHX in the presence or absence of TPCA-1 (0.5 μM) for 24 h, and cell viability was assessed (n = 3). (K) HL-7702 cells were treated with TNF+IFNγ in the presence or absence of rapamycin (100 nM) or TPCA-1 (0.5 μM) for 48 h, and cell viability was assessed (n = 3). (L) Mouse primary hepatocytes (WT and Raptor KO) were treated with Flag-TNF+CHX+z-VAD (TCZ) for the indicated times. Membrane-bound complex I was immunoprecipitated using anti-Flag antibody and representative proteins were analyzed by immunoblotting. (M) Mouse primary hepatocytes (WT, ATG9A KO, Raptor KO, and Raptor/ATG9A dKO) were treated with TCZ for the indicated times. Membrane-bound complex I was analyzed by immunoblotting. Asterisk indicates non-specific bands. (N) Mouse primary hepatocytes (WT and Raptor KO) were treated with TCZ in the presence or absence of TPCA-1 (0.5 μM), and representative proteins in complex I was analyzed. The data in all graphs are showed as mean ± SD. E and I show representative H&E staining from two independent experiments, G shows representative immunofluorescence from two independent experiments, F and J show representative immunoblots from three independent experiments. Statistical significance was determined using the log-rank (Mantel–Cox) test for survival curve comparisons (A, B, E, F, G), and two-way ANOVA for group comparisons (H, I, J): ****P < 0.0001, ***P < 0.001; ns, not significant.

**Supplementary Table 1.**
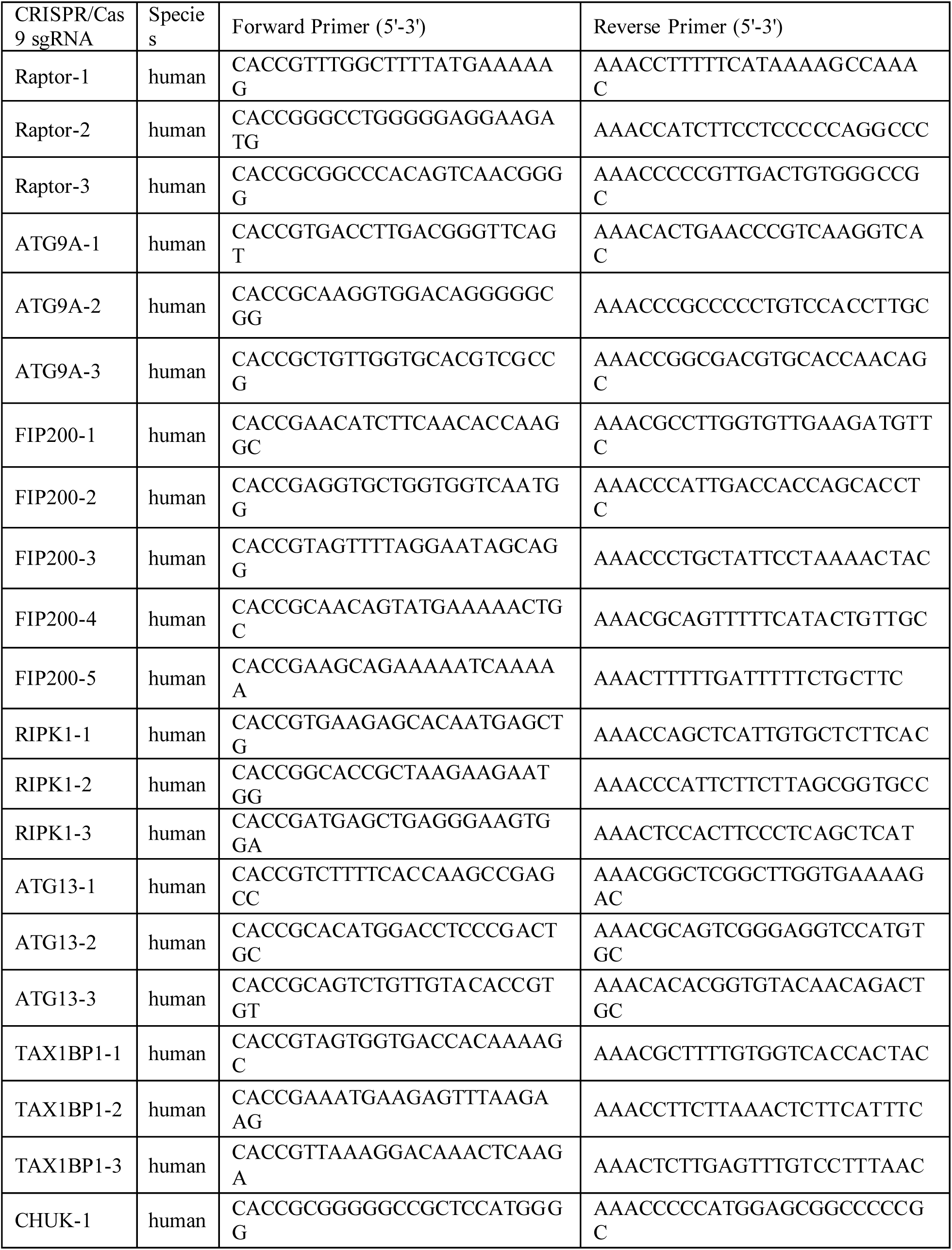
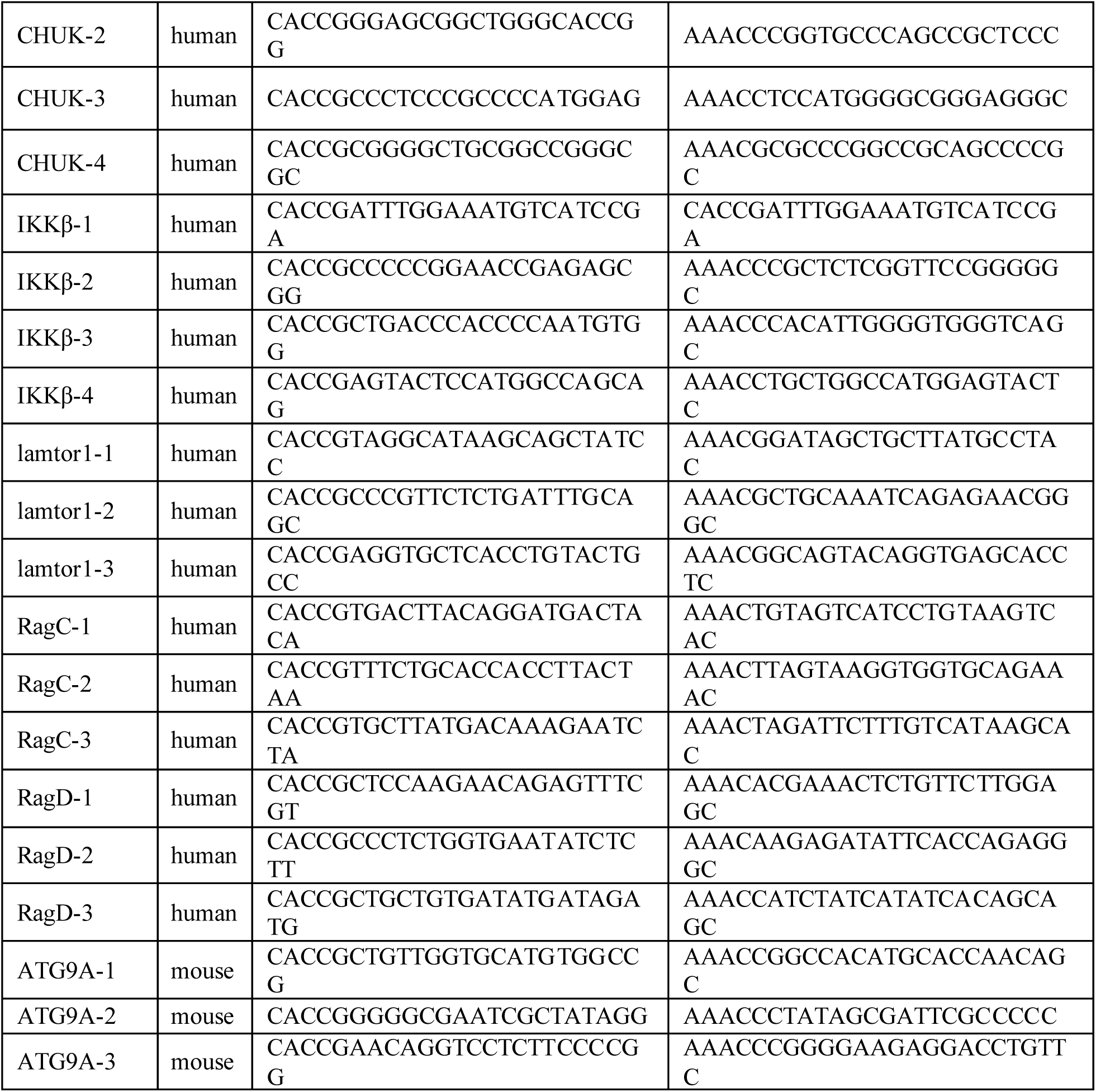
Primers used for sgRNA.

